# The LysR-type transcriptional regulator BsrA (PA2121) controls vital metabolic pathways in *Pseudomonas aeruginosa*

**DOI:** 10.1101/2021.06.08.447581

**Authors:** Magdalena Modrzejewska, Adam Kawalek, Aneta Agnieszka Bartosik

## Abstract

*Pseudomonas aeruginosa*, a facultative human pathogen causing nosocomial infections, has complex regulatory systems involving many transcriptional regulators. LTTR (LysR-Type Transcriptional Regulator) family proteins are involved in the regulation of various processes including stress responses, motility, virulence and amino acid metabolism. The aim of this study was to characterize the LysR-type protein BsrA (PA2121), previously described as a negative regulator of biofilm formation in *P. aeruginosa*. Genome wide identification of BsrA binding sites using ChIP-seq revealed 765 BsrA-bound regions in the *P. aeruginosa* PAO1161 genome, including 367 sites in intergenic regions. The motif T-N_11_-A was identified within sequences bound by BsrA. Transcriptomic analysis showed altered expression of 157 genes in response to BsrA excess, of which 35 had a BsrA binding site within their promoter regions, suggesting a direct influence of BsrA on the transcription of these genes. BsrA-repressed loci included genes encoding proteins engaged in key metabolic pathways such as the tricarboxylic acid cycle. The panel of loci possibly directly activated by BsrA, included genes involved in pili/fimbriae assembly as well as secretion and transport systems. In addition, DNA pull-down and regulatory analyses showed the involvement of PA2551, PA3398 and PA5189 in regulation of *bsrA* expression, indicating that this gene is part of an intricate regulatory network. Taken together, these findings reveal the existence of a BsrA regulon, which performs important functions in *P. aeruginosa*.

**IMPORTANCE:** This study shows that BsrA, a LysR-type transcriptional regulator from *P*. *aeruginosa*, previously identified as a repressor of biofilm synthesis, is part of an intricate global regulatory network. BsrA acts directly and/or indirectly as the repressor and/or activator of genes from vital metabolic pathways (e.g. pyruvate, acetate, tricarboxylic acid cycle), and is involved in control of transport functions and the formation of surface appendages. Expression of the *bsrA* gene is increased in the presence of antibiotics, which suggests its induction in response to stress, possibly reflecting the need to redirect metabolism under stressful conditions. This is particularly relevant for the treatment of infections caused by *P. aeruginosa*. In summary, the findings of this study demonstrate that the BsrA regulator performs important roles in carbon metabolism, biofilm formation and antibiotic resistance in *P. aeruginosa*.

## INTRODUCTION

Regulation of transcription is the principal mechanism controlling gene expression and the most economical way for a cell to respond to a rapidly changing environment. One of the largest groups of transcriptional regulators, with representatives in bacteria, archaea, and even eukaryotic organisms (1, 2), is the LysR Type Transcriptional Regulator (LTTR) family (3). Most LTTRs have two conserved and similarly organized functional domains (1, 4). The N-terminal DNA binding domain (DBD) with a winged helix-turn-helix motif mediates binding to cognate promoter sequences. The C-terminal effector binding domain (EBD), usually composed of two response subdomains (RD1 and RD2), is involved in ligand recognition and modulation of DBD activity (1, 3, 5). The conserved subdomain RD1 is also important for DNA interactions, whereas the more diverse RD2 contains an effector binding site (1, 6). LTTRs mediate signal-dependent and signal-independent transcriptional regulation of genes involved in numerous cellular processes, such as oxidative stress response, cell wall shape determination, quorum sensing, regulation of efflux pumps, secretion, motility, nitrogen fixation, virulence, cell division, metabolism and recognition of environmental stimuli and stresses (1).

The targets of LTTR regulation are often transcribed from a promoter that is very close to and may overlap that of a divergently transcribed regulator gene. In many cases, the LTTR positively regulates the target promoter in an effector-responsive manner, while negatively autoregulating its own promoter in the absence of an inducer (7–11). LTTRs can bind to target promoters in two conformations, depending on the presence of an effector. Ligand binding by the LTTR triggers a conformational change that permits binding to a DNA sequence involved in the regulation of its target gene. LTTRs may act as multimers, most frequently tetramers (12). Studies on several LTTRs have shown that the apoproteins can bind their promoters as tetramers, causing an extended DNase I footprint and a high-angle DNA bend, while the corresponding holoproteins produce a smaller footprint and lower DNA bend angle (1, 5). LTTRs usually bind to a sequence of approximately 50–60 bps, containing two distinct sites: a recognition-binding site or repression-binding site (RBS), encompassing the sequence T-N_11_-A (LTTR box), often located around position −65 relative to the start of transcription, and an activation-binding site (ABS) consisting of the −35 (ABS-35) and −10 (ABS-10) promoter regions (1, 5). In the absence of inducer, the LTTR tetramer binds to a RBS, but also with low affinity to the ABS-10 site, causing a bend in the DNA, leading to repression of the target gene by blocking availability of the −35 promoter region (13–15). The bent DNA is relaxed upon effector binding to the LTTR, leading to the formation of an active complex with RNA polymerase to initiate transcription. A ‘sliding dimer’ mechanism was proposed in which activation of the LTTR leads to a shift in the binding site from RBS/ABS-10 to RBS/ABS-35, releasing the −35 box for RNA polymerase recognition and subsequent gene expression (16, 17). Concomitantly, the autoregulatory properties of LTTRs are thought to be connected only with the dimeric form of the protein, not bound to the effector. The LTTR might bind to the RBS region of its own gene in a ligand-independent manner to regulate its expression (1).

One of the largest repertoires of LTTRs is encoded in the genome of *Pseudomonas aeruginosa*, an opportunistic human pathogen causing nosocomial infections including septicaemia, urinary tract infections, pneumonia, skin and wound infections (18–22). About 10% of all *P. aeruginosa* genes (usually around 6000) encode transcription factors. In the first sequenced *P. aeruginosa* genome of reference strain PAO1 (23), 113 genes are annotated as encoding LysR-type transcriptional regulators, but their functions remain largely unknown. *P. aeruginosa* LTTRs with known roles include PA0133 (BauR) (24), PA0739 (SdsB1) (25), PA1413 (26), PA1422 (GbuR) (27), PA1998 (DhcR) (28), PA2076 (OdsR) (29), PA2206 (30), PA2258 (PtxR) (31), PA2432 (BexR) (32), PA2838 (33), PA3225 (34), PA3587 (MetR) (35), PA3630 (GfnR) (36), PA4109 (AmpR) (37), PA4203 (38), PA5437 (PycR) (39), PA1003 (MvfR, also called PqsR) (40–42), PA5344 (OxyR) (43–45) and PA2492 (MexT) (46, 47). The membrane-associated multiple virulence factor regulator MvfR was shown to be necessary for *P. aeruginosa* virulence (40). MvfR positively regulates production of the *Pseudomonas* quinolone signal (PQS), one of three *P*. *aeruginosa* quorum sensing systems (48, 49), by controlling the *pqsABCDE* operon (50), as well as the *phnAB* genes involved in the biosynthesis of phenazine and anthranilic acid, a precursor of PQS (50, 51). Recent reports indicate that MvfR binds to dozens of loci across the *P. aeruginosa* genome at promoter regions, and within and outside the coding sequences of genes, recognizing different DNA binding motifs (41, 42), suggesting its involvement in the regulation of multiple genes. OxyR, another well characterized *P*. *aeruginosa* LTTR, is involved in the oxidative stress response, acting as a redox sensor (43). OxyR is activated by hydrogen peroxide (H_2_O_2_) and protects cells from toxic oxygen derivatives by stimulating the expression of the *katA*, *katB*, *ahpB* and *ahpCF* genes encoding catalases and alkyl hydroperoxide reductases (43, 52). It was recently shown that OxyR also regulates several other processes such as iron homeostasis, pyocyanin production and quorum sensing by binding to an AT rich motif (44, 45, 53). Another example of a *P. aeruginosa* LTTR with multiple roles is MexT (PA2492), an activator of the *mexEF*-*oprN* operon encoding a multidrug efflux pump involved in resistance to quinolones, chloramphenicol, trimethoprim and imipenem (46, 47, 54). Besides this handful of well-studied examples, the majority of LTTRs in this important pathogen remain uncharacterized.

Recently, a putative LTTR PA2121 was shown to negatively affect biofilm synthesis in the *P*. *aeruginosa* strain PAK and was therefore named biofilm synthesis repressor BsrA (55). It was shown, that the *bsrA* gene is regulated by the small regulatory protein SrpA during phage infection (56). SrpA is a key regulator controlling core cellular processes in *P. aeruginosa* PAK, including biofilm formation, and this factor binds to the motif TATC-N9-GATA identified within the *bsrA* promoter region.

In this study, we analyzed the role of BsrA in *P*. *aeruginosa* strain PAO1161, a derivative of PAO1 (57). In contrast to PAK, neither of these strains encodes *srpA* homologues. Our data indicate that the mode of BsrA action may differ in the strains PAK and PAO1161, because under the conditions tested, BsrA deficiency or overproduction had no influence on biofilm formation in PAO1161. Using RNA sequencing and chromatin immunoprecipitation we identified a BsrA regulon, which encompasses a gene encoding a key enzyme of the tricarboxylic acid cycle (TCA), a small RNA, as well as genes engaged in different cellular processes, some that are potentially involved in biofilm production. Using a DNA pull-down assay and regulatory experiments, we showed that other LysR-type regulators bind and regulate the *bsrA* promoter. Thus, BsrA is a part of an intricate regulatory network, that controls metabolic pathways during adaptation to a changing environment.

## RESULTS

### Impact of *bsrA* deficiency or overexpression on bacterial physiology

To analyze the role of BsrA in *P*. *aeruginosa*, a PAO1161 Δ*bsrA* mutant was constructed. This mutant strain did not display any significant differences in growth in LB or M9 medium, colony morphology, swimming or swarming, compared to the wild type (WT) parental strain PAO1161 (Fig. S1ABC). In parallel, the effect of *bsrA* overexpression was tested by linking the gene to an IPTG-inducible promoter in plasmid pMEB63 (*lacI^Q^-tac*p*-bsrA*). No effects of BsrA overproduction on bacterial growth were observed when IPTG concentrations of ≤ 0.25 mM were used (Fig. S1D), whereas 0.5 mM IPTG reduced the rate of growth significantly compared to cells carrying the empty vector (Fig. S1D).

As *bsrA* was initially identified as a repressor of biofilm synthesis, the formation of biofilms by the strains lacking or overproducing BsrA was examined. The absence of *bsrA* had no effect on the production of a biofilm by cells grown in either LB or M9 medium (Fig. 1A). Furthermore, the addition of arginine or a sub-inhibitory concentration of streptomycin to the growth medium, two compounds known to promote biofilm synthesis in *P*. *aeruginosa* (58, 59), resulted in comparable increases in biofilm formation in WT and Δ*bsrA* cells (Fig. 1A). Similarly, an excess of BsrA did not affect biofilm formation (Fig. 1B). These data suggested that BsrA may play an auxiliary or strain-specific role in biofilm formation in *P. aeruginosa*.

**Figure 1.**
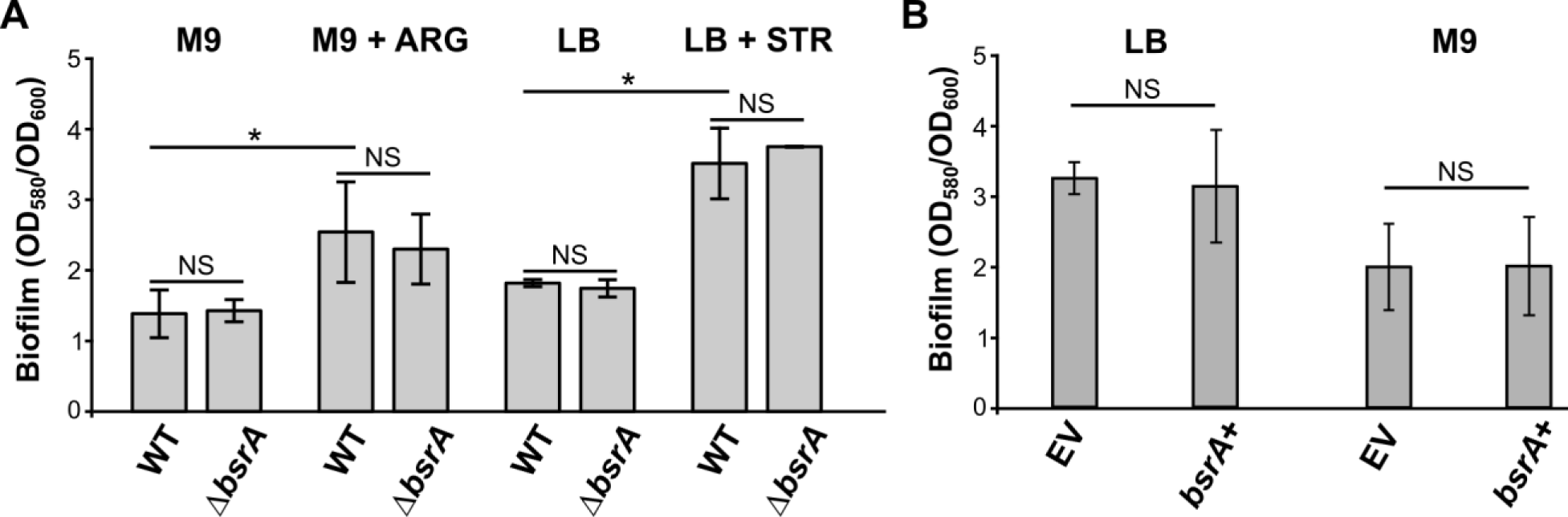
A lack or excess of BsrA does not affect biofilm formation by *Pseudomonas aeruginosa* PAO1161. Biofilm production in static cultures of (**A**) PAO1161 wild-type and the Δ*bsrA* strain grown in M9 medium supplemented with citrate as the carbon source (with or without arginine) or in LB medium (with or without 8 µg/ml streptomycin) for 48 h, and (**B**) the strain carrying pMEB63 (*lacI^Q^-tac*p*-bsrA)* overexpressing BsrA (*bsrA*+) and a control strain carrying empty vector (EV) pAMB9.37 (*lacI^Q^-tac*p), grown in medium supplemented with 0.05 mM IPTG for 72 h. OD_600_ values were measured and biofilm formation was assessed by staining with crystal violet, followed by measurement of OD_580_. Data represent the mean OD_580_/OD_600_ ratios ± SD from 5 biological replicates. * *p*-value <0.05 in a two-sided Student’s *t*-test. NS – not significant (*p*-value > 0.05).

### Identification of BsrA-regulated genes and binding sites for this transcriptional regulator in the *P*. *aeruginosa* genome

To identify genes that display BsrA-dependent expression we used RNA sequencing analysis (RNA-seq) to characterize the transcriptome of *bsrA*-overexpressing cells. In addition, we performed chromatin immunoprecipitation and sequencing analysis (ChIP-seq) to identify BsrA binding sites in the *P*. *aeruginosa* genome. The rationale behind an analysis of cells with BsrA in excess rather than the Δ*bsrA* mutant, was based on the following: 1) the relatively low level of *bsrA* expression under standard growth conditions (LB or M9 medium, data not shown); 2) the likelihood that an excess of BsrA might mimic the induced, activated state of the protein, and 3) the fact that the effector for this LTTR is unknown.

RNA-seq was performed using material isolated from cultures of the strains PAO1161 pMEB63 (*lacI^Q^-tac*p-*bsrA*, hereafter called BsrA+) and PAO1161 pAMB9.37 (*lacI^Q^-tac*p, empty vector - EV) grown in selective LB supplemented with 0.05 mM IPTG (Data set S1). Comparison of the BsrA+ and EV transcriptomes identified 157 loci with altered expression [fold change (FC) ≤ −2 or ≥ 2, FDR adjusted *p*-value ≤ 0.01] (Fig. 2A; Data set S2). The expression of 65 loci was down-regulated, while 92 loci displayed increased expression. For convenience, we use the *P. aeruginosa* PAO1 gene names throughout the manuscript, although the corresponding PAO1161 gene names are included in all tables. Functional classification of the identified loci, based on PseudoCAP (60), showed that the up-regulated genes were mostly involved in protein secretion/export systems, adaptation and protection as well as cell wall functions (Fig. 2B; Data set S2). Decreased expression was observed for several genes encoding proteins engaged in carbon compound metabolism and central intermediary metabolism. The most severely down-regulated genes were *PA3452* (*mqoA*), encoding a malate:quinone oxidoreductase from the TCA cycle and *PA0887* (*acsA*) encoding an acetyl-coenzyme A synthetase (61, 62), while the most highly up-regulated loci were the *mexXY* operon, encoding a multidrug efflux RND transporter (63–65), as well as genes encoding type VI secretion proteins (*PA1657*-*PA1671*) and transporters (*PA4192*-*PA4195*, *PA2202*, *PA2203*, *PA5024*) (Data set S2). The altered expression of selected loci in response to BsrA excess was confirmed using RT-qPCR analysis (data not shown).

**Figure 2.**
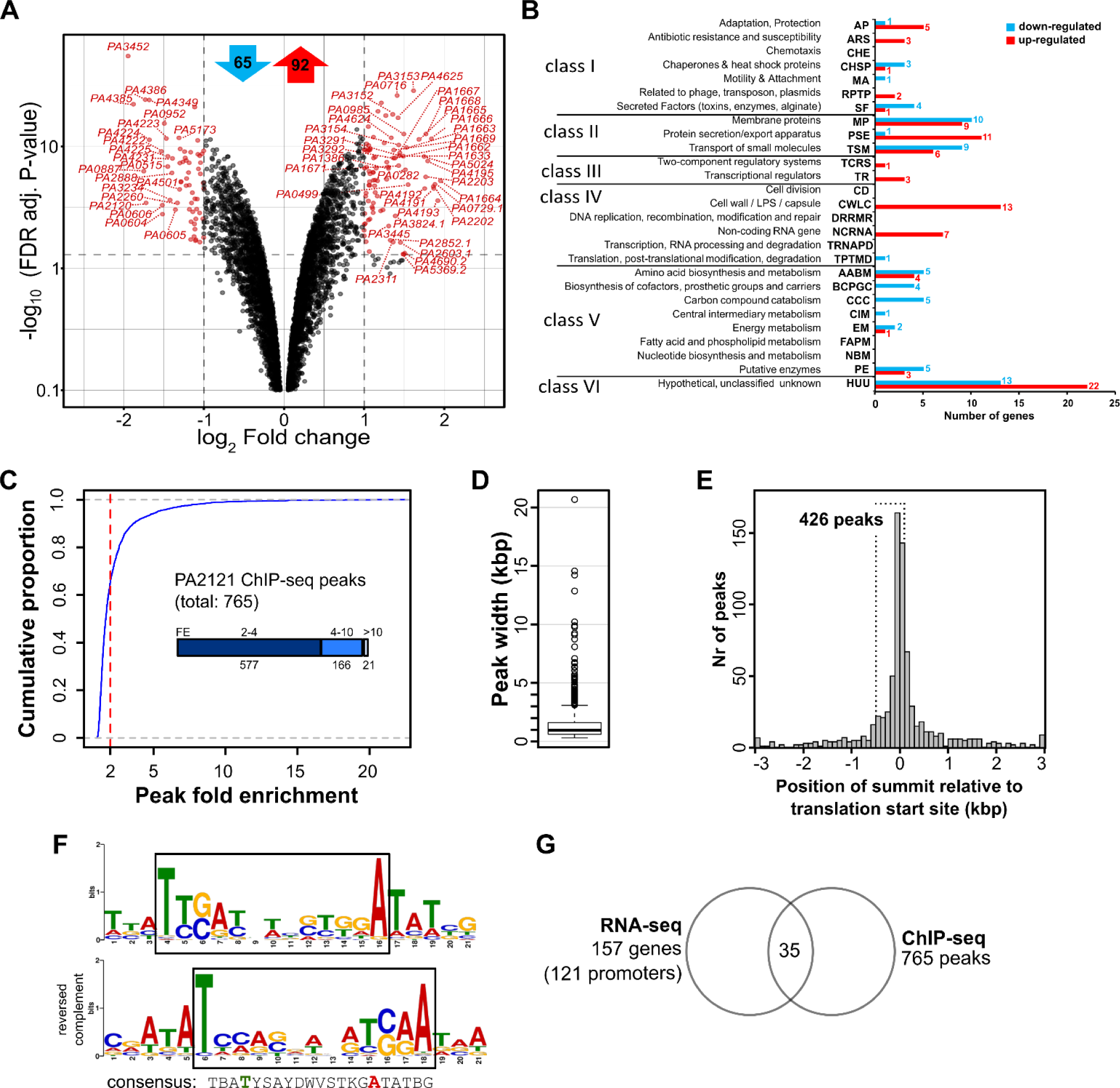
Identification of BsrA-dependent genes and binding sites for this transcriptional regulator in *P. aeruginosa*. Transcriptomes of PAO1161 cells carrying pMEB63 (*tac*p-*bsrA*, overexpressing BsrA - BsrA+) or pAMB9.37 (*tac*p, empty vector control - EV), grown under selection in LB supplemented with 0.05 mM IPTG were analyzed by RNA-sequencing. (**A**) Volcano plot of RNA-seq data comparing the transcriptomes of BsrA+ and EV cells. Differentially expressed genes (Fold change >2 or < −2, and FDR-corrected *p*-value ≤ 0.01) are indicated in red and the genes with the most significant changes in expression are named. For clarity, genes with a *p*-value < 0.1 are not shown. The numbers of up- and down-regulated loci are presented at the top in red and blue arrows, respectively. (**B**) Classification of loci with altered expression in response to BsrA excess according to PseudoCAP categories (60). When a gene was assigned to multiple categories, the most informative category was selected (in bold in Data set S2). The PseudoCAP categories were additionally grouped into six classes (103, 109). Red and blue bars correspond to the numbers of up- and down-regulated genes, respectively. (**C**) Identification of BsrA binding sites in the *P. aeruginosa* genome. Cells expressing BsrA-FLAG (or the control) were subjected to chromatin immunoprecipitation using anti-FLAG antibodies. Reads obtained by sequencing of the ChIP DNA were mapped onto the PAO1161 genome (57) and peaks were called using MACS2 with data from a mock-treated sample as the normalisation control. The chart represents the distribution of fold enrichment (FE) values for the detected peaks. A cut-off value of 2 is indicated by a red line. (**D**) Width distribution of BsrA ChIP-seq peaks. (**E**) Distribution of the distance between ChIP-seq peak summits and the nearest start codon. Bin width is 100 nt. Peaks with distances of > 3 kbp are grouped together in boundary bins. (**F**) Sequence logo of the BsrA binding motif obtained by MEME (66). The reverse complement of this logo and a proposed consensus sequence are presented below. B – C or G or T, Y – C or T, S – G or C, D – A or G or T, W – A or T, K – G or T. The LTTR box (T-N_11_-A) is framed in black. (**G**) Overlap between RNA-seq and ChIP-seq results. A gene was classified as likely to be directly regulated by BsrA if the ChIP-seq peak summit was located in the region −500 to +100 from its start codon (or the start codon of the corresponding operon).

To identify BsrA binding sites in the *P*. *aeruginosa* genome, ChIP-seq analysis was performed using an anti-FLAG antibody and Δ*bsrA* cells carrying plasmid pMEB99 (*tac*p-*bsrA*-*flag*), grown in selective LB supplemented with 0.05 mM IPTG. Addition of a FLAG-tag to the C-terminus of BsrA did not alter its ability to retard bacterial growth when overproduced (Fig. S2), indicating that the fusion protein is functional. As a background control for the ChIP procedure, the Δ*bsrA* strain carrying plasmid pABB28.1 (*tac*p*-flag*) was grown under the same conditions and samples were processed in parallel. Comparison of BsrA-FLAG ChIP samples with control samples, using a fold enrichment (FE) cut-off value of 2 (Fig. 2C) yielded 765 BsrA-FLAG ChIP-seq peaks (Data set S3). The majority of peaks exhibited an FE of between 2 and 4, although 166 had FE values of 4 to 10, and 21 had an FE of > 10 (Fig. 2C). The mean width of ChIP-seq peaks was < 1000 (twice the length of the DNA fragments used for ChIP), indicating BsrA binding to single or closely spaced binding site(s) (Fig. 2D). The summits of 367 peaks (48%) mapped to intergenic regions (Data set S3). A similar analysis of peak summit positions relative to the start codons of PAO1161 open reading frames (or the first genes in operons) showed that 426 peaks were located in the −500 to +100 regions, which suggests that the expression of these loci could be regulated by BsrA (Fig. 2E).

An extensive search for nucleotide motifs shared by sites bound by BsrA using MEME (66), showed the presence of a consensus sequence, resembling the T-N_11_-A motif (LTTR box) (Fig. 2F) proposed as the binding site of other LTTRs (1, 67, 68). These data indicated that BsrA has multiple binding sites in the *P. aeruginosa* genome, which suggests that this factor may function as a modulator of gene expression in regulatory networks.

### Genes under the direct control of BsrA

Interestingly, 35 of the 157 genes showing altered expression in response to a BsrA excess possessed a binding site for this transcriptional regulator within their promoter regions (Fig. 2G, Fig. 3ABC, Table 1). In addition, 55 BsrA peaks detected in coding regions were in the vicinity of genes that showed changes in expression level (fold change > 1.5) in RNA-seq analysis (Data set S3), but the mechanism by which BsrA could influence their expression requires further studies.

**Table 1.** Genes of *P. aeruginosa* likely to be regulated by BsrA, identified by ChIP-seq analysis. Loci with BsrA binding site(s) in the promoter regions preceding the genes and showing altered expression in response to a BsrA excess were considered to be directly regulated.

| Peak number | First gene of operon in PAO1161 (D3C65_) | First gene of operon in PAO1 | Position of summit relative to start codon | Gene in PAO1161 (D3C65_) | Gene in PAO1 | FC (fold change in RNA-seq) | FE (fold enrichment in ChIP-seq) | Gene description |
| --- | --- | --- | --- | --- | --- | --- | --- | --- |
| 188 | 07865 | PA3452 | -341 | 07865 | PA3452 | -3.86 | 6.05 | malate:quinone oxidoreductase |
| 79 | 03195 | PA0606 | -482 | 03195 | PA0606 | -2.87 | 2.73 | AgtD, ABC transporter permease |
| 534 | 21160 | PA0952 | -244 | 21160 | PA0952 | -2.82 | 2.13 | hypothetical protein |
| 751 | 29570 | PA5445 | -89 | 29570 | PA5445 | -2.46 | 2.47 | acetyl-CoA hydrolase/transferase family protein |
| 627 | 24675 | PA4542 | -54 | 24675 | PA4542 | -2.24 | 2.42 | ClpB, chaperone protein |
| 53 | 02095 | PA0396 | 12 | 02095 | PA0396 | -2.17 | 2.82 | PilT/PilU family type 4a pilus ATPase |
| 508 | 20310 | PA1112.1 | -56 | 20310 | PA1112.1 | -2.12 | 12.70 | non-coding RNA |
| 11 | 00575 | PA0105 | -198 | 00575 | PA0105 | -2.03 | 3.36 | CoxB, cytochrome c oxidase subunit II |
| 405 | 16285 | PA1874 | -216 | 16285 | PA1874 | 2.00 | 3.90 | Ig-like domain repeat protein |
| 188 | 07875 | PA3450 | -278 | 07875 | PA3450 | 2.03 | 6.05 | LsfA, 1-Cys peroxiredoxin |
| 372 | 14370 | PA2231 | -139 | 14330 | PA2239 | 2.04 | 4.77 | PsII, glycosyltransferase family 1 protein |
| 552 | 21955 | PA0805 | -28 | 21955 | PA0805 | 2.08 | 2.13 | hypothetical protein |
| 180 | 07670 | PA3488 | -305 | 07670 | PA3488 | 2.09 | 2.45 | hypothetical protein |
| 410 | 16485 | PA1838 | -7 | 16490 | PA1837 | 2.10 | 2.02 | DUF934 domain-containing protein |
| 348 | 13300 | PA2440 | -374 | 13295 | PA2441 | 2.10 | 2.45 | hypothetical protein |
| 389 | 15535 | PA2020 | -72 | 15535 | PA2020 | 2.11 | 2.50 | MexZ, transcriptional regulator |
| 652 | 25380 | PA4673.1 | 28 | 25380 | PA4673.1 | 2.15 | 7.15 | tRNA-Met |
| 189 | 07895 | PA3446 | -31 | 07895 | PA3446 | 2.16 | 5.54 | NADPH-dependent FMN reductase |
| 209 | 08850 | PA3266 | -21 | 08850 | PA3266 | 2.16 | 2.70 | CspA, cold-shock protein |
| 481 | 18935 | PA1372 | -193 | 18940 | PA1371 | 2.17 | 2.90 | DUF2290 domain-containing protein |
| 443 | 17455 | PA1656 | -253 | 17445 | PA1658 | 2.20 | 4.09 | TssC, type VI secretion system contractile sheath large subunit |
|  |  |  |  | 17450 | PA1657 | 2.20 |  | TssB, type VI secretion system contractile sheath small subunit |
| 309 | 12080 | PA2667 | -148 | 12080 | PA2667 | 2.22 | 2.95 | MvaU, H-NS family transcriptional regulator |
| 320 | 12420 | PA2602 | -74 | 12420 | PA2602 | 2.31 | 2.07 | 3-mercaptopropionate dioxygenase |
| 118 | 05080 | PA3981 | -46 | 5070 | PA3983 | 2.32 | 2.02 | HlyC/CorC family transporter |
| 646 | 25245 | PA4648 | -191 | 25250 | PA4649 | 2.00 | 6.09 | CupE2, Pilin subunit |
|  |  |  |  | 25245 | PA4648 | 2.43 |  | CupE1, Pilin subunit |
| 276 | 11045 | PA2852.1 | 64 | 11045 | PA2852.1 | 2.51 | 2.26 | tRNA-Ser |
| 643 | 25115 | PA4624 | -16 | 25115 | PA4624 | 2.53 | 2.12 | cyclic diguanylate-regulated TPS partner B, CdrB |
| 528 | 20980 | PA0985 | -471 | 20980 | PA0985 | 2.82 | 2.58 | pyocin S5 |
| 66 | 02635 | PA0499 | -89 | 02635 | PA0499 | 2.93 | 8.75 | probable pili assembly chaperone |
| 389 | 15540 | PA2019 | -94 | 15540 | PA2019 | 5.94 | 2.50 | MexX/AmrA family multidrug efflux RND transporter periplasmic adaptor |
|  |  |  |  | 15545 | PA2018 | 6.34 |  | MexY/AmrB family multidrug efflux RND transporter permease subunit |
|  |  |  |  | 15550 | - | 8.27 |  | transporter |
| 379 | 14975 | PA2121 | -8 | 14975 | PA2121 | 953.92 | 10.08 | LysR family transcriptional regulator |

Our analysis confirmed that BsrA might bind within the region preceding its own coding sequence (Fig. 3A). A BsrA binding site was also detected in the putative promoter of *PA3452* (*mqoA*): the gene showing the most severe down-regulation in the RNA-seq analysis (FC= −3.86) (Fig. 3B). Among the genes that might be directly regulated by BsrA, *PA1112*.*1*, encoding a small non-coding RNA (ncRNA) of unknown function (69), had a peak with the greatest fold enrichment (12.7) in the region preceding the structural gene (Fig. 3C).

**Figure 3.**
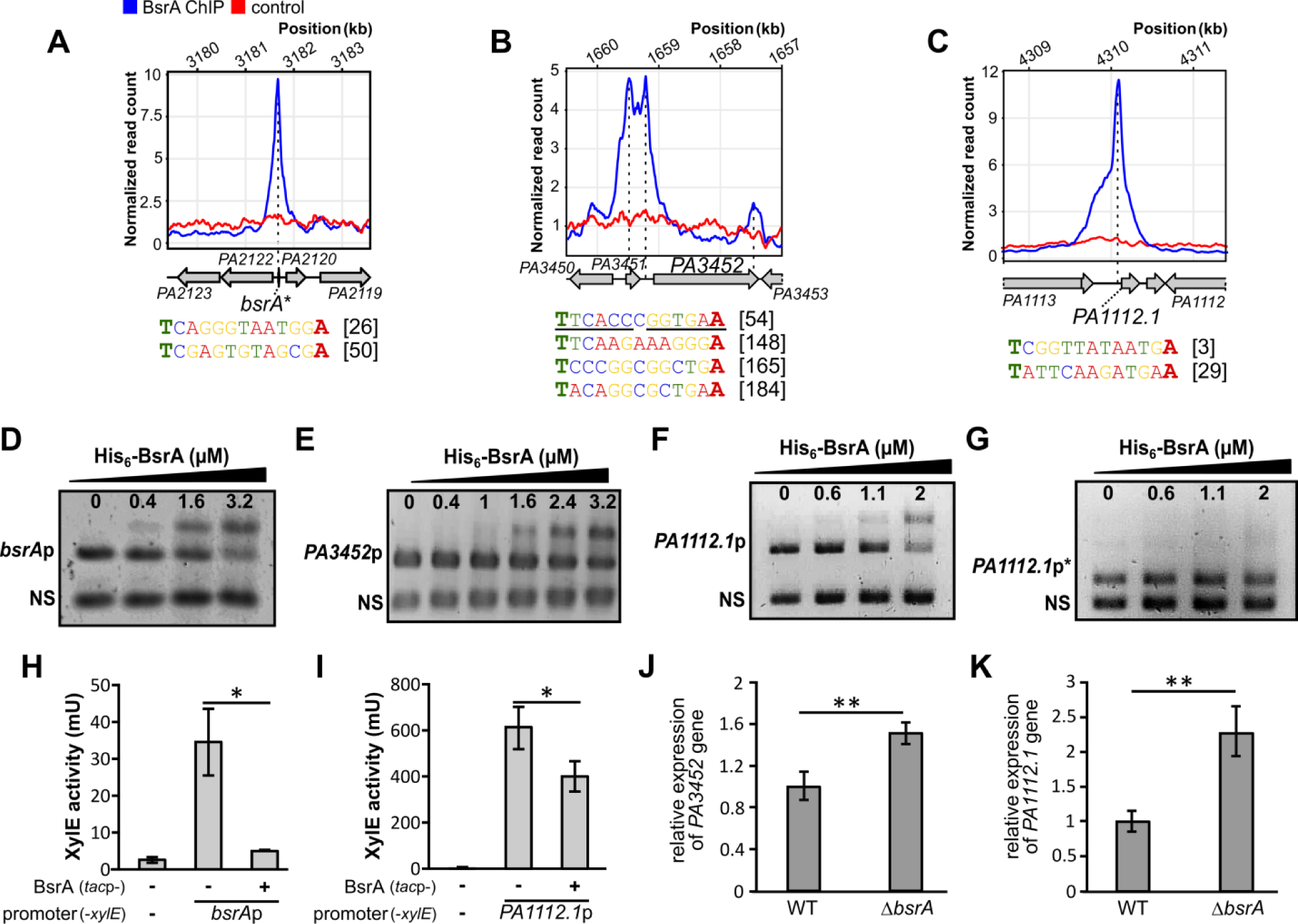
Direct regulation of target promoters by BsrA binding. ChIP-seq signal over the regions preceding the *bsrA* (**A**), *PA3452* (**B**) and *PA1112*.*1* (**C**) genes. The plots show normalized read counts, averaged for ChIP replicates, for the indicated positions in the PAO1161 (CP032126.1) genome. Genes are represented as arrows and the names of the PAO1 orthologues are shown for clarity. Sequences within the analyzed promoter fragments that correspond to the T-N_11_-A motif are presented below the plots, including their position relative to the start codon (underlined sequences indicate a pseudo-palindrome). EMSA analysis of His_6_-BsrA binding to regions preceding *bsrA* (**D**), *PA3452* (**E**) *PA1112.1* (**F**) and truncated *PA1112.1*p (lacking 71 bp containing the T-N_11_-A motif) (**G**). DNA fragments (0.1 µM) were incubated with the indicated amounts of His_6_-BsrA and complexes were separated by electrophoresis on 1.5% (**DEF**) or 2.5% (**G**) agarose gels stained with ethidium bromide. A 199-bp fragment of empty vector pCM132 (labeled as NS) was used as a control of binding specificity and a competitor DNA. XylE activity in *E*. *coli* DH5α double transformants carrying pMEB190 (*bsrA*p-*xylE*) (**H**) or pMEB232 (*PA1112.1*p-*xylE*) (**I**) plus pMEB63 (*lacI^Q^-tac*p-*bsrA*) for BsrA overproduction (+) or control plasmid pAMB9.37 (-). Strains were grown in selective LB. Data for cells carrying the promoter-less pPTOI (-*xylE*) and pAMB9.37 are shown as background controls. The data represent the means ±SD from three biological replicates. * indicates *p* < 0.05 in a Student’s two-tailed t-test. Relative expression (RT-qPCR) of *PA3452* (**J**) and *PA1112.1* (**K**) in WT and Δ*bsrA* cells from exponentially growing cultures (OD_600_ 0.2) normalized to the reference gene *rpsL*. ** indicates *p* < 0.01 in a Student’s two-sided t-test assuming equal variance.

To confirm the interactions of BsrA with putative promoters of these genes, we performed electrophoretic mobility shift assays (EMSA) using purified His_6_-BsrA and DNA fragments corresponding to the putative promoter regions of *bsrA*, *PA3452* and *PA1112*.*1*. Shifts of the promoter fragment DNA bands, but not of a non-specific competitor DNA were observed, indicating that His_6_-BsrA binds to these regions *in vitro* (Fig. 3DEF). To verify the importance of the LTTR box sequences in DNA binding by BsrA, version of the *PA1112.1* promoter fragment lacking the T-N_11_-A motif was tested in an EMSA. No BsrA binding to this shortened fragment (232 bp instead of 303 bp) could be detected (Fig. 3G).

To further examine the influence of BsrA on the expression of the three aforementioned genes, their promoter regions were cloned upstream of a promoter-less *xylE* gene in the vector pPTOI. The *bsrA* and *PA1112.1* promoters were active in the heterologous host *E*. *coli* DH5α, whereas no activity was observed for *PA3452*p (Fig. 3HI and data not shown). Expression of BsrA in cells carrying plasmids with *bsrA*p-*xylE* or *PA1112.1*p-*xylE* resulted in significantly reduced XylE activity in the corresponding cell extracts (Fig. 3HI). Moreover, RT-qPCR analysis of *PA3452* (*mqoA*) and *PA1112.1* transcript levels in *bsrA*-deficient cells showed increased expression of these two genes relative to WT cells, which supported the repressive effect of BsrA on the transcription of these genes (Fig. 3JK).

These data confirmed that BsrA binds to DNA fragments identified in ChIP-seq analysis and may regulate the activity of target promoters to influence gene expression. In addition, the T-N_11_-A nucleotide sequence, known as the LTTR box, present in the binding sites of most LTTRs (1, 67), is recognized by BsrA.

### Modulation of different cellular processes by BsrA

The RNA-seq results suggested that BsrA is engaged in modulating the activity of proteins mediating the conversion of malate to oxaloacetate in the TCA cycle by repressing the expression of the *PA3452* (*mqoA*) and *PA4640* (*mqoB*) genes (Data set S1). This is likely to influence subsequent steps of the cycle, e.g. the availability of oxaloacetate, its conversion to citrate using acetyl-CoA or the levels of acetyl-CoA generated via the pyruvate shunt (Fig. 4A). In addition, several genes that are putatively involved in the acetate transport (*PA3233*, *PA3234*) (70) and acetate pathways [*acsA* (*PA1562*), *acsB* (*PA1787*), *exaC* (*PA1984*)], encoding probable succinyl-CoA/acetate CoA-transferase (*PA5445*) (71), also showed reduced expression (FC >1.5) in response to BsrA (Fig. 4A; Table S1) (72, 73), suggesting the involvement of this LTTR in controlling acetate metabolism. We cultured the WT and Δ*bsrA* strains in minimal medium supplemented with citrate or acetate as the sole carbon source, but no visible effects on the kinetics of growth were observed (Fig. S1B). To test the effect of BsrA on acetate metabolism, the two strains were also cultured in medium containing a sub-inhibitory concentration of kanamycin, following the report of Meylan and co-workers, showing the effect of central carbon metabolite stimulation on aminoglycoside sensitivity in *P. aeruginosa* (74). The propagation of cells from overnight cultures in M9 medium containing 50 µg/mL kanamycin and acetate as the sole carbon source resulted in an increase in cfu/ml (relative to the starting point) of the *bsrA*-deficient mutant, while the cfu/ml value of the WT strain was not significantly changed (Fig. 4B). This effect was not observed when pyruvate and fumarate (compounds from different parts of the TCA cycle), or acetate plus fumarate, were used as the carbon source(s). Thus, the *P. aeruginosa* Δ*bsrA* mutant exhibited higher survival and/or fitness than the WT strain in the presence of kanamycin when grown in minimal medium supplemented with acetate as the sole carbon source, which confirmed the influence of BsrA on acetate metabolism.

**Figure 4.**
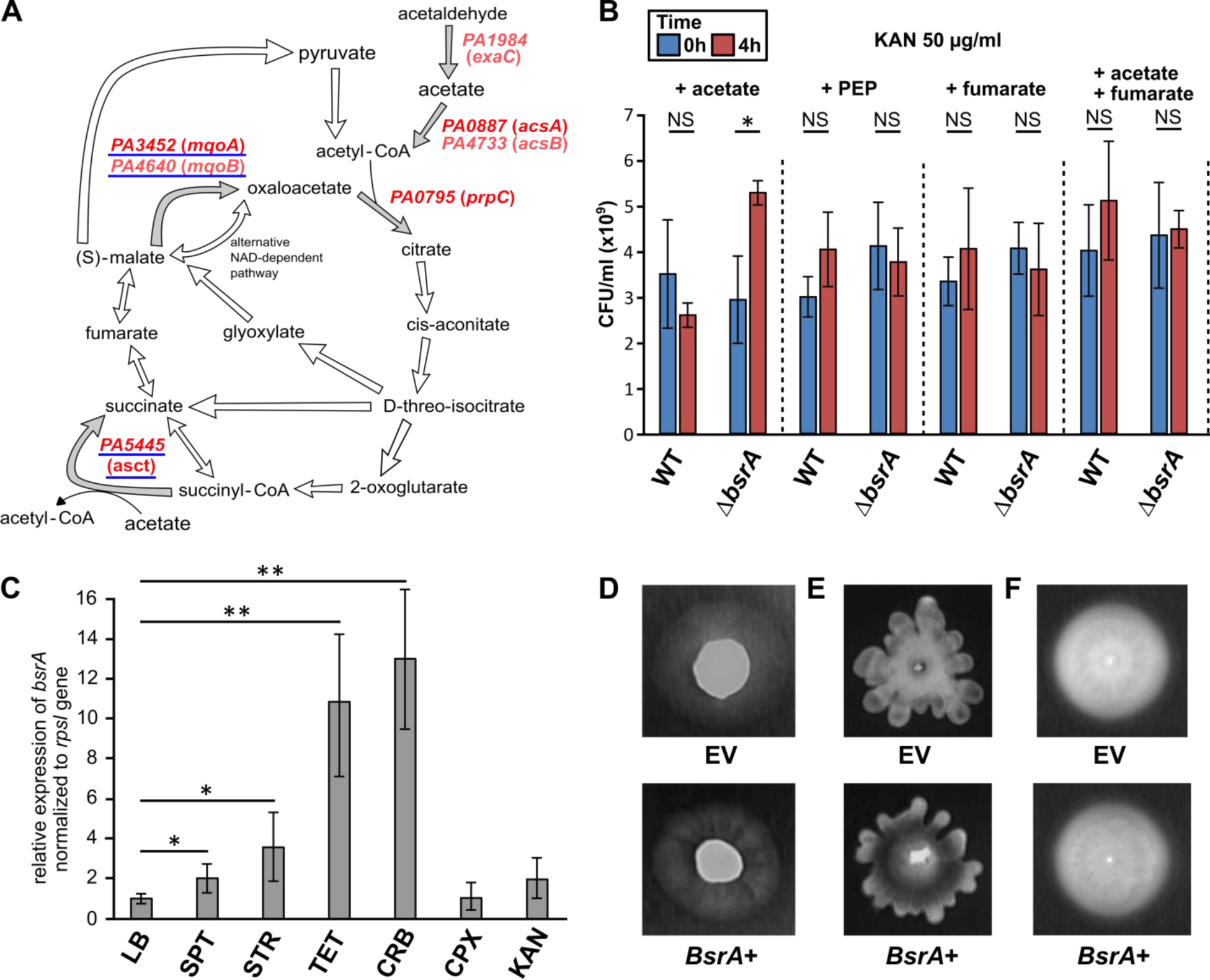
BsrA participates in the regulation of different processes in *P. aeruginosa* PAO1161. (**A**) Scheme of the TCA cycle (71, 73, 82, 96). Genes identified as affected by BsrA overproduction are indicated in red (dark red - FC > 2). Genes with BsrA binding sites in their promoters are underlined. (**B**) Viable cell density (cfu/ml) of overnight cultures of PAO1161 and the Δ*bsrA* mutant grown in M9 medium with kanamycin (50 µg/ml) and sodium acetate, phosphoenolpyruvate (PEP) or fumarate added as the sole carbon source, in amounts adjusted to maintain a total carbon concentration of 60 mM. * indicates a *p*-value of < 0.05 in Student’s *t*-test assuming equal variance. (**C**) Relative expression of *bsrA* in WT PAO1161 cells cultured in LB without antibiotic (LB) and with different classes of antibiotic added at sub-inhibitory concentrations: spectinomycin 128 µg/mL (SPT), streptomycin 4 µg/mL (STR), tetracycline 4 µg/mL (TET), carbenicillin 32 µg/mL (CRB), ciprofloxacin 0.06 µg/mL (CPX) and kanamycin 10 µg/mL (KAN). * and ** respectively indicate *p*-values of < 0.05 and < 0.01 in a Student’s two-sided t-test assuming equal variance. Twitching (**D**), swarming (**E**) and swimming (**F**) motility of strain PAO1161 carrying pMEB63 (overexpressing BsrA, *bsrA*+) or empty vector pAMB9.37 (control, EV) treated with 0.05 mM IPTG.

To test the effect of various antibiotics on *bsrA* expression, we performed RT-qPCR using RNA isolated from PAO1161 cultures grown in medium supplemented with sub-inhibitory concentrations of different antibiotics. This analysis showed no significant difference in *bsrA* expression upon the addition of kanamycin or ciprofloxacin compared with a negative control culture (Fig. 4C). Interestingly, the expression of *bsrA* was significantly increased in response to spectinomycin, streptomycin, tetracycline and carbenicillin (Fig. 4C), which indicates that *bsrA* is induced in response to specific antibiotics.

Our RNA-seq and ChIP-seq results also indicated increased expression of genes involved in fimbriae assembly (e.g. *PA0499*, *PA4648*-*PA4653*) in response to BsrA in excess. PA0499 is a periplasmic protein predicted to act as a chaperone assisting the assembly of appendages on the surface of the bacterium (75). *PA4648* is the first gene of the six-component *cupE* cluster encoding a so-called chaperone-usher pathway, the activation of which leads to the production and assembly of CupE fimbriae on the cell surface (76). These fimbriae are known to play a crucial role in biofilm development by *P. aeruginosa* and the *cupE* operon is specifically expressed in biofilm-forming cells (76). Since biofilm formation was unaffected in both the Δ*bsrA* and BsrA+ strains (Fig. 1), we checked whether *bsrA* overexpression had any effect on swimming, twitching or swarming motilities (77–79). BsrA+ cells showed differences in twitching (involves pili) and swarming, as demonstrated by the presence of clear radiating motility zones (“lines”) spreading from the centre of bacterial colonies, that were not observed in the control strain. This may reflect possible changes in radial expansion of the colony, which could be related to enhanced appendage production in BsrA+ cells (Fig. 4DE). No effect on swimming was observed (Fig. 4F), indicating that BsrA overproduction does not have a general negative effect on the motility of cells grown on plates. Thus, BsrA appears to be involved in the regulation of swarming and twitching motilities, and possibly attachment to surfaces, the first stage in biofilm formation.

Taken together, these results demonstrated the participation of BsrA in a number of diverse cellular processes including the modulation of cellular metabolism in response to growth conditions and the control of appendage formation leading to altered motility of *P. aeruginosa* cells.

### BsrA is under the control of other transcriptional regulators in *P. aeruginosa*

The findings of a previous study (55) and our data showed that the expression of *bsrA* is subject to autoregulation. To identify other proteins that can modulate *bsrA* transcription and in consequence the level of BsrA, we used a *bsrA* promoter fragment as bait in a DNA pull-down assay with *P. aeruginosa* PAO1161 cell extracts. The proteins bound to *bsrA*p were then characterized by mass spectrometry analysis. Altogether, 39 proteins were identified as being able to bind to *bsrA*p, but not to a control DNA fragment Data set S4). Importantly, BsrA was identified among the proteins with the highest scores, providing a positive control for this approach and confirming the autoregulatory properties of the protein. Six other proteins were identified with high scores for binding to *bsrA*p in two independently tested samples (eluates): PA2551, PA3587 (MetR), PA4902, PA4462 (RpoN), PA5189 and PA3398. Interestingly, five of these proteins are classified as LysR family transcriptional regulators, whereas PA4462 (RpoN) is a σ^54^ factor interacting with RNA polymerase (80). It is known that σ^54^ factors direct RNAP to conserved −12 (TGC) and −24 (GG) elements, and similar regions (TGA at position −12 and GG at position −24) are present in the *bsrA*p.

To determine whether the proteins identified in pull-down analysis can indeed affect the activity of the *bsrA* promoter, the *PA2551*, *PA3398*, *PA3587*, *PA4902* and *PA5189* genes were cloned under the control of *tac*p in vector pAMB9.37 and expressed in cells carrying plasmid pMEB190 (*bsrA*p-*xylE*). Measurements of XylE activity in cell extracts of the double transformants showed that the expression of *PA3587* and *PA4902* did not significantly influence *bsrA*p activity under the tested conditions (Fig. 5A). Notably expression of *PA2551*, *PA3398* or *PA5189* resulted in major decreases in XylE activity, suggesting that these proteins act directly as repressors of the *bsrA* gene. Interestingly, ChIP-seq analysis revealed strong binding of BsrA upstream of the *PA2551*, *PA3398* and *PA5189* genes (Fig. 5B), but not *PA3587* or *PA4902* (data not shown).

**Figure 5.**
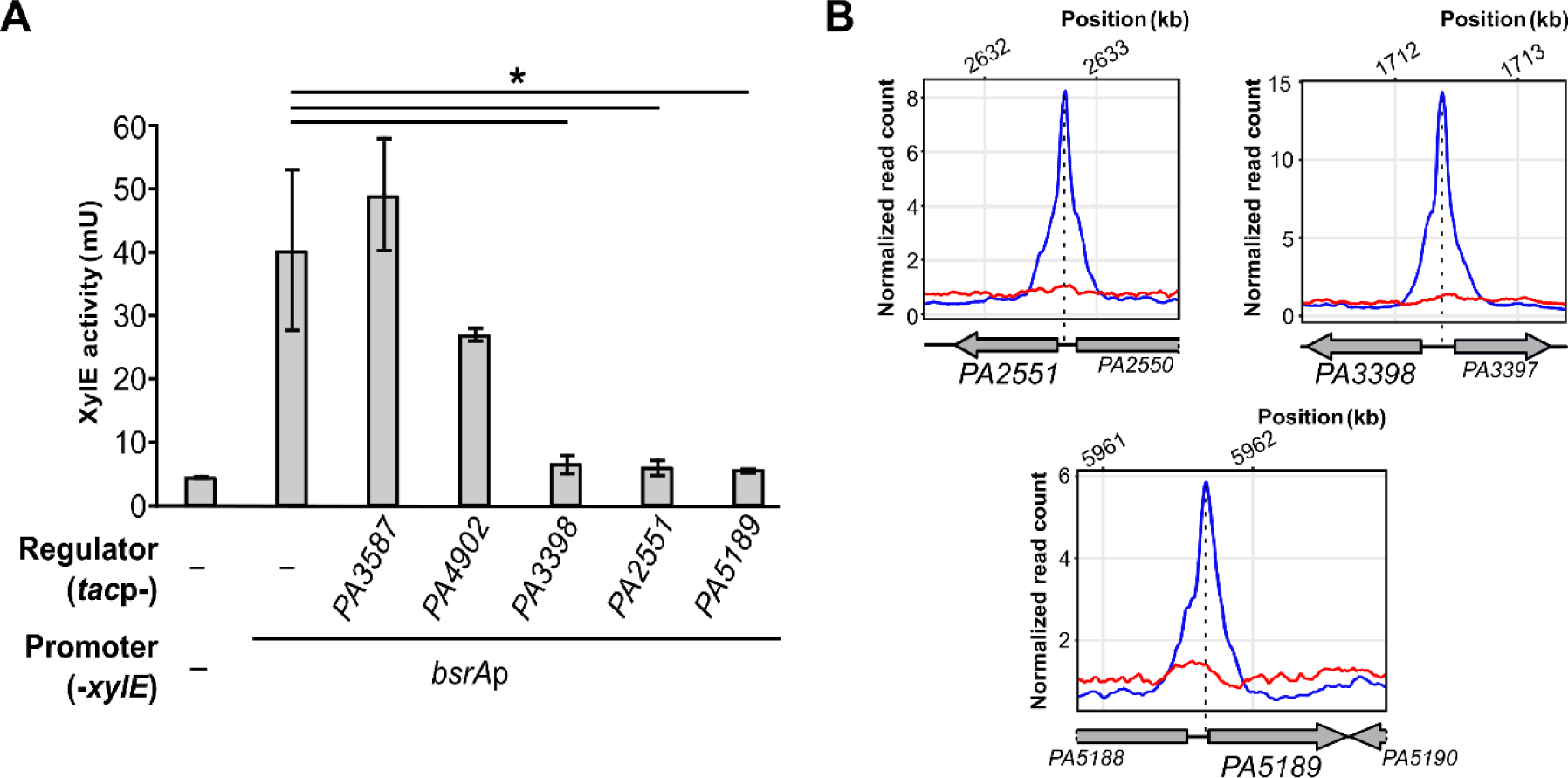
Transcriptional regulators PA2551, PA3398 and PA5189 control expression of *bsrA*. (**A**) XylE activity in double transformants of *E*. *coli* DH5α carrying promoter-less pPTOI or pMEB190 (*bsrA*p-*xylE*) plus vectors expressing the indicated genes under *tac* promoter control. Cells were grown in selective LB supplemented with 0.05 mM IPTG. Data represent the means ±SD from three biological replicates. * indicates *p* < 0.05 in a Student’s paired two-tailed *t*-test. (**B**) ChIP-seq signals over regions preceding the *PA2551*, *PA3398* and *PA5189* genes encoding regulators repressing *bsrA* expression. The plots show normalized read counts, averaged for replicates, for the indicated positions in the PAO1161 (CP032126.1) genome.

These results showed that BsrA is part of an intricate regulatory network involving mutual regulation between BsrA and other LysR-type transcription factors.

## DISCUSSION

In this study, we performed a functional analysis of the LysR-type transcriptional regulator BsrA (PA2121) from *P*. *aeruginosa*, previously described as a repressor of biofilm synthesis (55).

Transcriptional analysis of a strain overproducing BsrA revealed the greatest changes in gene expression for loci encoding enzymes engaged in carbon metabolism (mainly down-regulated) and for loci predicted or known to be involved in processes connected with transport, biofilm and type VI secretion systems (up-regulated). In a *P. aeruginosa* PAK mutant with disrupted *bsrA*, increased biofilm synthesis was observed (55), while the PAO1161 Δ*bsrA* mutant constructed in this study did not show significant changes in biofilm formation (Fig. 1). This difference could be related to the presence of the SrpA protein in the PAK strain, which is not encoded in the genome of PAO1161 (or PAO1). Among its other functions, SrpA directly regulates expression of the *bsrA* gene by binding to its promoter (56). Our data suggested that another mechanism is responsible for regulating biofilm production, possibly involving BsrA-mediated activation of genes such as *PA0499* or *PA4648*, that have been connected with the formation of biofilms (75, 76). The generation of biofilm structures is strictly linked to metabolic activity, inhibited in the cells of the mature biofilm matrix, but increased during early biofilm development (81). The role of BsrA in biofilm formation might be related to the modulation of these processes. The relationship between SrpA and BsrA in biofilm formation requires further study.

Our data suggested that BsrA is involved in the repression of metabolic functions by direct or indirect down-regulation of genes engaged in pyruvate metabolism and the TCA cycle (Fig. 4A; Fig. 6). The most highly repressed gene, directly controlled by BsrA, is *mqoA* (*PA3452*) encoding a putative malate:quinone oxidoreductase (MQO), a FAD-dependent enzyme involved in the conversion of malate to oxaloacetate. The gene encoding the second *P. aeruginosa* MQO, *mqoB* (*PA4640*), was also subject to BsrA-mediated regulation, although to a lesser extent. The presence of *mqoB* is necessary for the growth of cells on acetate and ethanol as sole carbon sources (82). Under these conditions, one of the primary functions of MQOs is to replenish the oxaloacetate pool in the TCA cycle to allow further assimilation of acetyl-CoA and permit TCA operation to provide intermediates for biosynthetic processes and respiration (82). Both MQOs are produced by cells grown under standard aerobic conditions, but levels of MqoB are higher than those of MqoA (33, 73, 82). The precise role of MqoA in *P. aeruginosa* awaits elucidation. Bacterial MQOs have previously been characterized in *E. coli* and *Corynebacterium glutamicum* as the principal enzymes catalysing the oxidation of malate (61, 83). In *Pseudomonas putida* the *mqo-2* gene, encoding a malate:quinone oxidoreductase 2, is under the control of Crc, the global regulator of carbon catabolite repression (CCR) (84). The assimilation of energetically favourable carbon sources is the main bacterial strategy employed to optimize metabolism and growth. Crc protein together with the RNA chaperone translational repressor Hfq and small RNA(s) comprise the CCR regulatory system in pseudomonads (85, 86). In *P. aeruginosa*, a specific sRNA named CrcZ has been identified as an antagonist of Crc and Hfq. CrcZ binds to the Crc and Hfq proteins, trapping and sequestering them. The expression of *crcZ* is under the control of a two-component system CbrA/CbrB, which reacts to carbon source availability (85, 87, 88). It is clear that a multilevel regulatory network involving sRNAs plays an important role in metabolic regulation in pseudomonads, which is interesting in light of our identification of a sRNA (*PA1112.1*) as a target of BsrA regulation.

**Figure 6.**
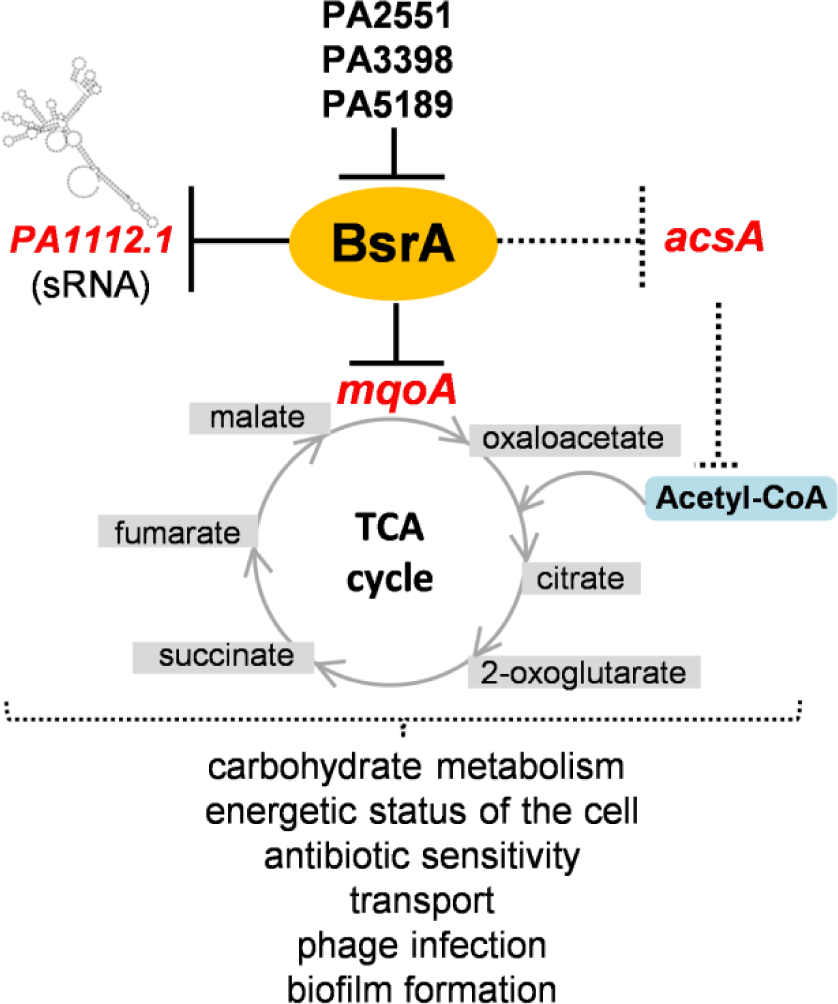
The BsrA regulatory network in *P. aeruginosa* and its impact on bacterial physiology. A black solid line indicates direct repression by this transcriptional regulator; a dotted line indicates direct and/or indirect involvement of BsrA in the control of gene expression and downstream processes.

Our analysis of the phenotype of the *bsrA*-deficient mutant demonstrated its increased fitness in the presence of kanamycin compared to the WT strain under specific conditions. It was previously recognized that the efficacy of aminoglycoside antibiotics depends on metabolic stimuli (74, 89, 90). As an aminoglycoside, kanamycin acts by inhibiting protein synthesis through binding to the 30S subunit of the bacterial ribosome. Killing of bacterial cells by kanamycin depends on proton-motive force (PMF), which is required for the uptake of the antibiotic (89). PMF is related to NADH level, which is dependent on metabolism. Therefore, the cellular metabolic state modulates the uptake and/or efficacy of the antibiotic (90). Although adaptation to antibiotics is thought to be controlled at the transcriptional level by the induction of stress responses, several reports have indicated that there is a relationship between a high concentration of certain endogenous metabolites and the level of bacterial resistance (91–94). We found that the Δ*bsrA* mutant displayed better adaptation to kanamycin under conditions of acetate supplementation and it may be speculated that this is due to altered drug uptake due to changes in PMF generation, a process connected with the TCA cycle and cellular respiration (89, 90, 95). Growth on acetate requires the activity of the glyoxylate shunt which supplies cells with malate and oxaloacetate (Fig. 4A). It might be connected with reoxidization of the NADH excess generated by the TCA cycle during growth on acetate and the need to coordinate the composition of the electron transport chain at the level of the terminal oxidases, e.g. the proton pumping NADH dehydrogenase I or Nqr (73). The Δ*bsrA* mutant had an increased level of the transcript of malate dehydrogenase *mqoA* (Fig. 3J) and probably those encoding several other enzymes from the TCA cycle and acetate metabolism. The lack of repression of TCA cycle enzymes or genes involved in acetate metabolism in the Δ*bsrA* mutant in comparison with the WT strain may provide some advantage during growth on acetate in the presence of kanamycin and adaptation to the stress caused by this antibiotic.

Kanamycin sensitivity was examined in cells grown on other carbon sources, but a significant difference in antibiotic adaptation of the Δ*bsrA* mutant was only observed with acetate supplementation. The main reason for this may be the stage at which particular carbon compounds enter the TCA cycle, as shown by Dolan and co-workers (73). These authors presented so-called “carbon fluxes” leading to metabolic and transcriptomic changes caused by growth on acetate or glycerol. We speculate that the lack of BsrA leads to elevated TCA cycle flux connected with metabolism remodelling when acetate is the sole carbon source.

An interesting gene belonging to the BsrA regulon, potentially connected with TCA cycle remodelling, is *PA5445.* This gene putatively encodes succinyl-CoA/acetate CoA-transferase, an enzyme engaged in the conversion of succinyl-CoA and acetate to succinate and acetyl-CoA, which could modulate the TCA cycle and confer some advantage during growth on acetate. PA5445 displays almost 50% identity to AarC from *Acetobacter aceti*, a bacterium utilizing a specialized TCA cycle (71). In this bacterium AarC-mediated conversion of succinyl-CoA to succinate replaces the action of typical succinyl-CoA synthetases (SucC, SucD) (71, 96). This modification is connected with enhanced tolerance to low pH and acetate, produced by *Acetobacte*r during fermentation. Many bacteria including *P. aeruginosa* possess homologues of *aarC* (*asct*) in addition to the *sucC* and *sucD* genes, which suggests the existence of an alternative pathway in the TCA cycle, possibly conferring some advantage connected with acetate metabolism (96).

Similarly to *mqoA*p, the promoter region of *PA5445* possesses few potential BsrA binding sites (matching the consensus in Fig. 2F), with one putative site (TTCGACCTTGGTA) overlapping the predicted −10 promoter region and located very close to a BsrA ChIP-seq peak summit. This suggests that BsrA may regulate genes encoding components of metabolic pathways and can mediate metabolism remodelling, which could lead to increased fitness of the Δ*bsrA* mutant in the presence of kanamycin.

Interestingly, *bsrA* (*PA2121*) was identified as one of a panel of genes containing mutations in *P. aeruginosa* cystic fibrosis isolates, which may have been selected during adaptation and evolution to promote survival during infection of the lungs of these patients (97–99). In addition, Kong and co-workers (100), using a *luxCDABE*-based random promoter library of *P. aeruginosa* PAO1, identified *PA2121* (*bsrA*) as one of 45 genes that perform a role in long-term survival and thus may be involved in chronic infections of the human body.

BsrA binds to numerous sites in the *P. aeruginosa* genome, yet it only had a limited influence on the regulation of gene expression under the conditions tested (Data set S3). The majority of BsrA binding sites contain the LTTR box, composed of the sequence T-N_11_-A, but besides this element there is a low level of sequence conservation. It was not possible to define a more specific binding motif, which suggests the involvement of other factors in mediating BsrA binding to DNA. This observation highlights the potentially broad role of BsrA in modulating gene expression in *P. aeruginosa*, with the possible involvement of other regulatory proteins that associate with sequences adjacent to BsrA binding sites under specific growth conditions. The nature of the signal to which BsrA responds and the precise role of this factor, require further study.

Recently, high-throughput SELEX analysis has been used to define the preferred binding motifs of 53 *P*. *aeruginosa* LysR-type transcriptional regulators (101). Most of these LTTRs display dimeric binding to cognate sequences. The recognised binding sites are mostly palindromic or have partial dyad symmetry and range in length from 12 to 24 base pairs. Sequence conservation is highest within the flanking regions, that usually display dyad symmetry, whereas there is often very low sequence conservation inside the motif. In most of the binding sites the LTTR-box (T-N11-A, T-N10-A or T-N9-A) can be identified as part of the sequence creating dyad symmetry. The motif preferentially recognized by BsrA was identified as NAGTAGACNNGTCTACTN; however, no such sequence was found in the genomes of PAO1 or PAO1161 and no highly similar sequences were present in the regions identified using ChIP-seq analysis. FIMO analysis (102) using 200-bp sequences encompassing the BsrA peak summits identified only 5 sequences with a *p*-value of < 0.0001 resembling the proposed motif [peaks 682, 194, 367, 276, 157 (Data set S3)] or 56 sequences when a *p*-value cut-off of 0.001 was used. The preferential BsrA binding site motif identified in our analysis is more generic, but is recognizable as an LTTR box characteristic for LysR-type regulators, and better explains the presence of multiple LTTR binding sites within the promoters of cognate genes.

LTTRs usually bind to promoters of target genes upstream from the transcription start site. Among the tested promoter regions of BsrA regulated genes, i.e. *bsrA*, *mqoA* and *PA1112.1*, two to four T-N_11_-A motifs, closely resembling the BsrA binding site (Fig. 2F) were identified (Fig. 3A-C). These are located at positions from 3 to 184 bp from the start codon of these down-regulated gene, so that BsrA binding to these sites might reduce RNA polymerase access to the core promoter sequences (−10, −35). To specifically recognize and bind cognate DNA, LTTRs use highly conserved interactions between amino acids and nucleotide bases as well as numerous less conserved secondary interactions (7, 68). One site, often called the recognition binding site, consists of a T-N_11_-A motif with imperfect dyad symmetry. It is believed that interaction with this site anchors the LTTR to the DNA and is often involved in repression, including autoregulation. LTTRs are known to bind to longer sequences (50-60 bps) containing a so-called activation binding site, and these interactions are usually driven by the presence of a specific ligand or co-factor, which is bound by the LTTR. In addition, LTTRs bind with higher or lower affinity to their binding sites depending on the presence or absence of its inducer or ligand, which modulates interaction with DNA. Conformational flexibility of the created LTTR multimers (usually tetramers), causes DNA bending or relaxation, which regulates the repression or activation state of the regulator (13). Conformational changes may also permit transient contacts of the regulator with DNA sequences flanking the T-N_11_-A motif, which might also be affected by occupation by other DNA-interacting factors. The availability of the regulator in the cell, the possibility of creating monomers or multimers to exert a regulatory effect on target promoters, as well as the dynamic order in which different binding events take place, which determines the appropriate regulatory response, could provide further levels of control. Our pull-down results highlighted the existence of an intricate regulatory network engaging in possible crosstalk, cooperation and/or interconnection between different transcriptional regulators exerting an influence on *bsrA* expression and further on its targets. Thus, different factors control LTTR interactions with DNA, providing specificity of recognition and correct timing of this action.

Based on the presented results, we propose a model of the regulatory network engaging BsrA in *P. aeruginosa* and its impact on bacterial physiology (Fig. 6). BsrA acts as the repressor of genes involved in carbohydrate metabolism (*mqoA*, *acsA*) influencing the TCA cycle, the availability of acetyl-CoA and overall cellular metabolism. In addition, BsrA regulates the transcription of the uncharacterized sRNA *PA1112.1*, which is possibly involved in post-transcriptional regulation of gene expression. Interestingly, besides autoregulation, the *bsrA* gene is under the control of other LTTRs of *P. aeruginosa* (PA2551, PA3398 and PA5189) indicating the ability to fine tune BsrA action in the cell. This multilevel regulatory network plays a role in controlling carbohydrate metabolism (TCA cycle, acetate and pyruvate metabolism) and thus the energetic status of the cell, which has implications for other functions such as cellular transport, the response to antibiotic, phage infection, biofilm formation, virulence and overall survival strategies. In line with this model, the induction of *bsrA* expression was observed in the presence of antibiotics and also in *parA* and *parB* mutants characterized by growth retardation and defects in chromosome distribution (103), which suggests the release of *bsrA* expression in response to stress and the need to redirect metabolism to cope with adverse conditions, that might be manifested by a slowdown of bacterial growth.

## MATERIALS AND METHODS

### Bacterial strains, plasmids and growth experiments

Bacterial strains used and constructed in this study (listed in Table S1) were grown in LB or on LB-agar at 37°C, and in M9 minimal medium supplemented with sodium citrate (0.25%) or sodium acetate (20 mM) as the carbon source, with leucine (10 mM) added in the case of *P. aeruginosa* PAO1161 *leu^−^* strains. For the selection of plasmids in *E. coli*, media were supplemented with 10 µg/ml chloramphenicol, 50 µg/ml kanamycin or benzyl penicillin at a final concentration of 150 µg/ml in liquid medium or 300 µg/ml in agar plates. For *P. aeruginosa* strains, carbenicillin (300 µg/ml), rifampicin (300 µg/ml), kanamycin (250 µg/ml in liquid medium; 500 µg/ml in plates) and chloramphenicol (75 µg/ml in liquid medium; 150 µg/ml in plates) were applied as required.

For growth experiments, liquid media were inoculated with strains propagated on plates. These cultures were grown overnight with shaking at 37°C, diluted 1:100 in fresh medium and then incubation was continued. Bacterial growth was monitored by the measurement of optical density at 600 nm (OD_600_) at 1 hour interval. Competent *E*. *coli* cells were prepared by treatment with CaCl_2_ and transformation was performed according to a standard procedure (104). Competent *P*. *aeruginosa* cells were prepared as described previously (105).

All plasmids used and constructed in this study are described in Table S1.

A *P. aeruginosa* PAO1161 Δ*bsrA* mutant was obtained by allele exchange (106). Competent cells of *E. coli* S17-1 were transformed with plasmid pMEB14 (a derivative of suicide vector pAKE600) to create the donor strain, and WT *P. aeruginosa* PAO1161 Rif^R^ was used as the recipient. The allele exchange procedure was performed as described previously (106, 107). Verification of the obtained mutant strain was performed by PCR using primer pair #4/#7 (Table S2).

### Motility assays

Motility assays were performed as described previously (79), supplementing the swimming, swarming, and twitching media, if necessary, with chloramphenicol (150 µg/ml) and IPTG (0.05 mM). To standardize the assays, all plates contained the same volume of the medium.

### RNA isolation, RNA-seq and RT-qPCR

Total RNA was isolated from three independent replicate samples of *P. aeruginosa* PAO1161 overexpressing the *bsrA* gene as well as the control strain carrying the empty vector or *P. aeruginosa* PAO1161 WT and the Δ*bsrA* strain. RNA isolation and analysis were performed as described in Text S1.

### Chromatin immunoprecipitation with sequencing

ChIP was performed according to the procedure of Kawalek et al. (108) with some modifications, as described in Text S1.

### Protein purification

*E. coli* BL21(DE3) transformed with pMEB10 encoding a His_6_-BsrA fusion protein was grown to exponential phase in autoinduction LB medium (Foremedium) containing 1% (v/v) glycerol and 0.5% (w/v) NaCl. The cells were harvested by centrifugation, resuspended in phosphate buffer (50 mM sodium phosphate, pH 8.0) supplemented with lysozyme (1 mg/ml), PMSF (1 mM) and benzonase nuclease (250 U, Sigma), then sonicated. His_6_-BsrA was purified from the cell lysate by chromatography on Ni-agarose columns (Protino Ni-TED 1000, Macherey-Nagel) with 300 mM imidazole in phosphate buffer used for elution. The purification procedure was monitored by SDS-PAGE using a Pharmacia PHAST gel system. Fractions containing the purified protein were dialyzed overnight in Tris buffer containing 5% (v/v) glycerol and stored in small aliquots at −80°C.

### *In vitro* protein-DNA interactions

The electrophoretic mobility shift assay (EMSA) was performed to determine the ability of purified BsrA to bind to selected promoter regions of *P. aeruginosa* genes *in vitro*, as described in Text S1.

### Regulatory experiments with promoter-*xylE* fusions in *E. coli*

*E. coli* DH5α double transformants carrying pPT01 derivatives with the promoter regions of selected *P. aeruginosa* genes fused to the *xylE* reporter gene plus pAMB9.37 (*lacI^Q^*-*tac*p) derivatives expressing the tested proteins were assayed for catechol 2,3-oxygenase activity (the product of *xylE*) as described in Text S1.

### Tests of kanamycin sensitivity

The effect of kanamycin on PAO1161 cells was tested using the carbon source screening procedure (74, 89) described in Text S1.

### DNA pull-down assay

Pull-down analysis was performed as described previously (108) with modifications summarized in Text S1.

### Data availability

The raw RNA-seq and ChIP-seq data supporting the results of this article were deposited in the NCBI’s Gene Expression Omnibus (GEO) database (http://www.ncbi.nlm.nih.gov/geo/) under GEO Series accession numbers GSE163234 and GSE163233 (for release after manuscript acceptance).

## Supporting information

Data set S1

Data set S2

Data set S3

Data set S4

## SUPPLEMENTAL MATERIALS

**Text S1.** Materials and methods.

**Figure S1** Selected diagrams and charts presenting the phenotypic analysis of *P. aeruginosa* PAO1161 WT, Δ*bsrA* mutant, and the strain overproducing BsrA.

**A.** Growth curves of the *P. aeruginosa* PAO1161 *bsrA* mutant and WT strains in LB and M9+leucine+citrate at 37°C (*leu^-^* strains).
**B.** Growth curves of the *P. aeruginosa* PAO1161 *bsrA* mutant and WT strains in M9+citrate and M9+acetate at 37°C (*leu*^+^ strains).
**C.** Selected pictures of swimming and swarming assays.
**D.** Growth curves of *P. aeruginosa* PAO1161 strains carrying pMEB63 (*lacI^Q^*-*tac*p-*bsrA*; BsrA overproducer) in L-broth with gradient (0-0.5 mM) of IPTG inducer.

**Figure S2** Comparison of the impact of *bsrA* and *bsrA*-*flag* overexpression on the growth of cells in culture.

**A.** Growth curves of *P. aeruginosa* PAO1161 strain carrying pMEB99 expressing *bsrA*-*flag* fusion grown in L-broth with gradient (0-0.5 mM) of IPTG inducer.
**B.** Growth curves of *P. aeruginosa* PAO1161 *ΔbsrA* strains carrying pABB28.1 (*tacp-flag*; F-EV) and pMEB99 (*lacI^Q^*-*tac*p*-bsrA*-*flag*; BsrA-F) in L-broth with 0.5 mM IPTG (BsrA-F+; F-EV+) or without induction (BsrA-F; F-EV).

**Data set S1** Full RNA-seq data for BsrA+ and EV transcriptomes. Genes identified only in PAO1161 strain but not in PAO1 are described as “not annotated (NA)”.

**Data set S2** Results of RNA-seq analysis. List of 157 genes with altered expression identified by comparison of the transcriptomes of the BsrA+ and EV strains [fold change (FC) ≤ −2 or ≥ 2, FDR ≤ 0.05]. The PseudoCap categories in the bold text were the most informative and were used as the gene information presented in Fig. 2B. Genes identified only in strain PAO1161 but not in PAO1 are described as “not annotated (NA)”.

**Data set S3** Results of ChIP-seq analysis. 765 BsrA-FLAG ChIP-seq peaks with a fold enrichment (FE) cut-off value of ≥ 2 obtained by the comparison of BsrA-FLAG ChIP samples with negative control samples. Genes identified only in PAO1161 strain but not in PAO1 are described as “not annotated (NA)”.

**Data set S4** Proteins interacting with *bsrA*p in a pull-down assay, identified by mass spectrometry analysis. Only proteins binding *bsrA*p but not to the control fragment are shown.

**Table S1** Bacterial strains and plasmids used and constructed in this study.

**Table S2** List of primers used in this study.

## ACKNOWLEDGEMENTS

We thank Jan Gawor, Karolina Zuchniewicz and Robert Gromadka from the Laboratory of DNA Sequencing and Oligonucleotide Synthesis, IBB PAS, Warsaw, Poland for performing RNA and DNA sequencing. We thank the Laboratory of Mass Spectrometry of IBB PAS, Warsaw, Poland for the analysis of pull-down samples. This work was supported by the National Science Centre in Poland (grant 2015/18/E/NZ2/00675).

