## Supplementary material for "The LysR-type transcriptional regulator BsrA (PA2121) controls vital metabolic pathways in *Pseudomonas aeruginosa*": Table S1

**Table S1** Bacterial strains and plasmids used and constructed in this study.

| **strain** | **description** | **reference** |
| --- | --- | --- |
| ***Escherichia coli*** |  |  |
| DH5α | F^–^ Φ80*lacZ*Δ*M15* Δ(*lacZYA-argF*) *U169 recA1 endA1 hsdR17* (r_K–_, m_K+_) *phoA supE44 λ thi-1 gyrA96 relA1* | (1) |
| S17-1 | *pro* Δ*hsdR hsdM*^+^ *recA* Tp^R^ Sm^R^ ΩRP4-Tc::Mu Kn::Tn7 | (2) |
| BL21 | F^−^ *ompT hsdS*_B_ (r_B_^−^m_B_^−^) *gal dcm* (λ DE3) | Novagen |
| ***Pseudomonas aeruginosa*** |  |  |
| PAO1161 | PAO1161 Rif^R^ *leu*^-^, r^-^, m^-^ | (3) |
| PAO1161 | PAO1161 Rif^R^ r^-^, m^-^ | (4) |
| PAO1161 Δ*bsrA* | PAO1161 Rif^R^ with deleted gene *bsrA* (*PA2121*), allele exchange with the use of pMEB14 | This study |
| **name of the plasmid** | **description** | **reference** |
| pBBR1-MCS-1 | Cm^R^, IncA/C broad-host-range cloning vector, *lacZ*α–MCS, *mob*, T7p, T3p | (5) |
| pAMB9.37 | pBBR1-MCS-1 derivative with *lacI^Q^ tac*p, expression vector | (6) |
| pABB28.1 | pBBR1-MCS-1 derivative with *lacI^Q^* *tac*p-*flag*, expression vector | (7) |
| pMEB1 | pAMB9.37 derivative with modified *mcs* (added PstI, SmaI, BglII, SalI restriction sites) | This study |
| pAKE600 | Ap^R^, *ori*_MB1_, *oriT*_RK2_, *sacB*, suicide vector | (8) |
| pBGS18 | Km^R^, *ori*_MB1_, cloning vector | (9) |
| pCM132 | Km^R^, *oriC*_ColE1_, *oriV*_IncP_, *oriT*, *traJ*’, *trfA*, broad-host-range vector, promoter-less *lacZ* reporter gene | (10) |
| pET28a(+) | Km^R^, *ori*_MB1_, T7p, *lacO*, His_6_-tag, T7 tag, expression vector | Novagen |
| pPTO1 | Km^R^, *oriV*_pSC101_, promoter-less *xylE* cassette | (11) |
| pET28mod | Km^R^, *ori*_MB1_, T7p, *lacO*, His_6_-tag, modified to remove T7 tag | (12) |
| pKAB240 | Ap^R^, *ori*_MB1_, pUC19 derivative with His_6_-*mcs* (MunI, HindIII, NotI, XhoI, BamHI)-*flag* | (13) |
| pMEB5 | pAKE600 derivative with upstream region of *bsrA* gene inserted as EcoRI-HindIII fragment amplified with the use of #4 and #5 primers | This study |
| pMEB9 | pBGS18 derivative with downstream region of *bsrA* gene inserted as HindIII-BamHI fragment amplified with the use of #6 and #7 primers | This study |
| pMEB10 | pET28mod derivative, containing *bsrA* gene inserted as EcoRI-SacI fragment amplified with the use of #1 and #2 primers to obtain His_6_-*bsrA* fusion | This study |
| pMEB14 | pAKE600 derivative with fused upstream and downstream region of *bsrA* gene; downstream region from pMEB9 inserted into pMEB5 to create Δ*bsrA* allele | This study |
| pMEB47 | pCM132 derivative with 328 bps fragment containing intergenic region upstream of *bsrA* amplified with the use of #8 and #9 primers fused with *lacZ* to obtain transcriptional fusion | This study |
| pMEB63 | pAMB9.37 derivative with inserted EcoRI-XhoI fragment containing *bsrA* gene (cloned from pMEB10) under control of *tac*p promoter | This study |
| pMEB98 | pKAB240 derivative with *bsrA* gene fused with flag-tag (*bsrA*-*flag*); *bsrA* gene without stop codon amplified with the use of #1 and #3 primers inserted as EcoRI-XhoI fragment | This study |
| pMEB99 | pAMB9.37 derivative with inserted EcoRI-SalI fragment from pMEB98 containing *bsrA*-*flag* | This study |
| pMEB187 | pCM132 derivative with 348 bps fragment containing intergenic region upstream of *PA3452* amplified with the use of #10 and #11 primers fused with *lacZ* to obtain transcriptional fusion | This study |
| pMEB190 | pPTO1 derivative with 328 bps fragment containing intergenic region upstream of *bsrA* amplified with the use of #8 and #9 primers fused with *xylE* to obtain transcriptional fusion | This study |
| pMEB230 | pAMB9.37 derivative with *PA2551* gene amplified with the use of #18 and #19 primers, inserted as EcoRI-XhoI under control of *tac*p promoter | This study |
| pMEB231 | pAMB9.37 derivative with *metR* (*PA3587*) gene amplified with the use of #20 and #21 primers, inserted as EcoRI-XhoI under control of *tac*p promoter | This study |
| pMEB232 | pPTO1 derivative with 306 bps fragment containing intergenic region upstream of *PA1112.1* amplified with the use of #12 and #13 primers fused with *xylE* to obtain transcriptional fusion | This study |
| pMEB236 | pMEB1 derivative with *PA3398* gene amplified with the use of #22 and #23 primers, inserted as EcoRI-SalI under control of *tac*p promoter | This study |
| pMEB237 | pMEB1 derivative with *PA4902* gene amplified with the use of #24 and #25 primers, inserted as EcoRI-SmaI under control of *tac*p promoter | This study |
| pMEB238 | pAMB9.37 derivative with *PA5189* gene amplified with the use of #26 and #27 primers, inserted as EcoRI-XhoI under control of *tac*p promoter | This study |
