## Supplementary material for "The LysR-type transcriptional regulator BsrA (PA2121) controls vital metabolic pathways in *Pseudomonas aeruginosa*": Table S2

**Table S2** List of primers used in this study.

| **Primers used to amplify DNA fragments for cloning** | | |
| --- | --- | --- |
| nr | name | sequence 5**´**-3**´** |
| #1 | **2121ESF** | GC GAATTC ATGAAGCTGAACTTGCAGCAG |
| #2 | **2121ESR** | GC GAGCTC AGTGACGGAAGACTGATCCTC |
| #3 | **2121EX** | CG CTCGAG GCCCTGTTCGATCTGCAC |
| #4 | **2121EHuF** | GC GAATTC CACCGTCAGCCATTCCAGCAG |
| #5 | **2121EHuR** | GC AAGCTT TCATCACAGCTTCATGAATAATCCTC |
| #6 | **2121HBdF** | GC AAGCTT TGAGTCATAGGTTGCGCGCG |
| #7 | **2121HBdR** | GC GGATCC CCTCGAAGACCGTGAGCAC |
| #8 | **p2121EBF** | GC GAATTC GCATGC CAGTTGAAGAACACCAGC |
| #9 | **p2121EBR** | GC GGATCC CATGAATAATCCTCATTC |
| #10 | **p3452F** | GC GAATTC GCATGC CATAAGTTCGGTGAATGGGC |
| #11 | **p3452R** | GC GGATCC CGCTAGAAAACGTGGAGAAG |
| #12 | **p20325F** | GC GGTACC GCATGC CCCGGAGAACTTTCTCGCATC |
| #13 | **p20325R** | GC GGATCC ATCCGGGGACCTGTCTCTGCA |
| #14 | **BADNsiIF** | ACGGATGGCCTTTATGCATTTCTACAAACT |
| #15 | **CM132RCy5** | Cy5 - CTTCCACAGTAGTTCACCACC |
| #16 | **CM132RBiot** | Biotin - CGTCAGTAACTTCCACAGTAG |
| #17 | **CM132pF** | GTGAACGCTCTCCTGAGTAG |
| #18 | **2551EXf** | CG GAATTC ATGAACCTGAACAAAGTCGA |
| #19 | **2551EXr** | AT CTCGAG CTATCCGTCGTTTCCGCA |
| #20 | **3587EXf** | CG GAATTC ATGCTCGAACTTCGCCACC |
| #21 | **3587EXr** | CG CTCGAG TACCTACCGTTCGGCCTTCA |
| #22 | **3398ESlf** | CG GAATTC ATGAAATTCACCCTCCGCC |
| #23 | **3398ESr** | CT GTCGAC CTATACCTCGTACCGCCAG |
| #24 | **4902ESmf** | CG GAATTC ATGAGTCCGATCGATCTC |
| #25 | **4902ESr** | AT CCCGGG GTGTTGCCGCTGCTCGGT |
| #26 | **5189EXf** | CG GAATTC ATGGTGAATATCCAAACCTTC |
| #27 | **5189EXr** | AT CTCGAG GGAAGAACTGGTCATTCG |
| **Primers used in qPCR analysis** | | |
| #28 | **rpsLF** | CTCGGCACTGCGTAAGGTAT |
| #29 | **rpsLR** | TGTGCTCTTGCAGGTTGTGA |
| #30 | **pa3452qF** | AAGGCCTTCAAGGACAAGGT |
| #31 | **pa3452qR** | TCAGCTCGATGTCGTTGTTC |
| #32 | **20325qF** | GGCGAGATCCAGAAGATCG |
| #33 | **20325qR** | AGCGCTTCTTGCTGATGGT |
| #34 | **pa2121qF** | GCAGCACGTCTCGATCCT |
| #35 | **pa2121qR** | CGGGATGGTCAGGTAGTGAG |
