## Supplementary material for "The LysR-type transcriptional regulator BsrA (PA2121) controls vital metabolic pathways in *Pseudomonas aeruginosa*": Text S1

**Text S1.** Materials and methods

*P. aeruginosa* PAO1161 Δ*bsrA* mutant construction

To investigate the role of the BsrA protein in *P. aeruginosa*, a *bsrA* null mutant was constructed in *P. aeruginosa* PAO1161 strain. The WT and Δ*bsrA* strains were tested in several growth conditions showing no visible significant defects on bacterial growth under tested conditions (Fig. S1A). At the beginning of the research, the *leu^-^* PAO1161 strains were tested, but the need for leucine supplementation in minimal medium prevented studies on single carbon sources. Then the obtained prototrophic PAO1161 *leu^+^* strain (1) was used (Fig. S1B).

RNA isolation, RNA-seq, bioinformatic analysis of RNA-seq data

Strains were obtained by transformation of WT cells with pMEB63 or pAMB9.37 plasmids (Table S1). Transformants were selected on LB plates supplemented with 150 μg/ml chloramphenicol and were verified by isolation of plasmid DNA and its digestion. After overnight growth, each culture of *P. aeruginosa* PAO1161 carrying pMEB63 or pAMB9.37 vector, was diluted 1:100 into fresh LB medium supplemented with 75 μg/ml chloramphenicol and 0.05 mM IPTG as inducer. 2 ml of cultures which reached optical density 0.4-0.6 at 600 nm were mixed with 4 ml of RNAprotect Bacteria Reagent (Qiagen). RNA was isolated with an RNeasy mini kit (Qiagen) according to the manufacturer’s protocol. Total RNA was digested with DNase (TURBO DNA-free Kit, Ambion) to eliminate genomic DNA.

Sequencing of RNA was performed in the Laboratory of DNA Sequencing and Oligonucleotide Synthesis of Institute of Biochemistry and Biophysics Polish Academy of Sciences in Warsaw. Ribosomal RNA was depleted using RiboZero Bacteria kit (Illumina). Obtained mRNA was used for cDNA library construction using KAPA Stranded RNASeq kit. Library was further quality checked on 1% agarose gel and concentration was measured using qPCR KAPA Library Quantification Kit. Libraries were sequenced using standard Illumina protocols. Reads were quality-checked and filtered using FASTP version 0.20.0 (<https://github.com/OpenGene/fastp>) (2). Reads were mapped to *P*. *aeruginosa* PAO11161 genome (CP032126.1) using Bowtie2 version 2.3.4.3 (3) using default settings. The number of reads mapping to individual genes was counted using FeatureCounts v 2.0.1 (part of the Subread [http://subread.sourceforge.net](http://subread.sourceforge.net/)) with the -s2 option (4). Differential expression analysis was conducted using edgeR ver 3.28.0 (5). Raw data are available in the NCBI‘s Gene Expression Omnibus (GEO) database (http://www.ncbi.nlm.nih.gov/geo/) under accession number [GSE16323](https://www.ncbi.nlm.nih.gov/geo/query/acc.cgi?acc=GSE163233)4.

RT-qPCR

For RT-qPCR analyses, RNA from samples collected from three independent replicates of *P. aeruginosa* PAO1161 WT and Δ*bsrA* strain was isolated with a Total RNA Mini kit (A&A Biotechnology, Poland) according to the manufacturer’s protocol using 2 ml of cultures. cDNA was synthesised from total RNA with the use of Maxima First Strand cDNA Synthesis Kit (ThermoFisher Scientific). qPCR was performed with Hot FIREPol EvaGreen qPCR Mix Plus (Solis Biodyne) in the LightCycler 480 (Roche). The same amount of cDNA (~50 ng) was added to each mixture and reaction was performed in a total volume of 18 µl. Specific qPCR primers were used to amplify the reference *rpsL* gene (primers #28/#29) and target genes *PA3452* and *PA1112.1* (primers #30/#31 and #32/#33, respectively) (Table S2). Changes in gene expression between WT (wild type) and the Δ*bsrA* mutant were calculated with the use of Pfaffl method (6). Three biological replicates with three technical replicates were tested for each gene. RNA isolated the same way as described above from *P. aeruginosa* PAO1161 WT cultured in LB and LB with the addition of antibiotics such: spectinomycin 128 µg/ml (SPT), streptomycin 4 µg/ml (STR), tetracycline 4 µg/ml (TET), carbenicillin 32 µg/ml (CRB), ciprofloxacin 0.06 µg/ml (CPX) and kanamycin 10 µg/ml (KAN) till OD_600nm_ reached about 0.5 was used to obtain cDNA and perform RT-qPCR with the use of pair of primers amplifying the fragment of *bsrA* gene (#34/#35) and *rpsL* used as the reference gene (#28/#29).

Chromatin immunoprecipitation with sequencing

*P. aeruginosa* PAO1161 Δ*bsrA* strain overexpressing *bsrA* fused with FLAG-tag at C-end as well as control strain carrying empty vector pABB28.1 were obtained by conjugation and selection on LB agar plates supplemented with 300 µg/ml rifampicin and 150 µg/ml chloramphenicol. Four independent overnight cultures from each strain were inoculated and diluted 1:100 in LB medium supplemented with 75 µg/ml chloramphenicol and 0.05 mM IPTG to induce the expression of *bsrA*-*flag*. Bacteria were grown until reaching the exponential phase with OD_600_ about 0.5. The further procedure was performed as described earlier until the step of sonication (7). Lysate after sonication was thawed on ice and 150 μL of each strain variant was incubated with 20 μL of magnetic beads coupled with protein A (Dynabeads Protein A, Invitrogen, 10001D) previously separated from original suspension using Magnetic Separation Stand. The pre-clearing step was performed for 1 hour at 4^o^C with rotation of the mixtures. 50 μL of magnetic beads, separated from the suspension as above, was then mixed with anti-FLAG mouse polyclonal antibodies (1-10 μg per reaction, 3 and 6 μL in this procedure) [DYKDDDDK Tag polyclonal antibodies; PA1-985B; Invitrogen; Thermofisher Scientific; 1 mg/ml] diluted in 200 μL of PBS with 0.05% Tween-20. Mixtures of magnetic beads and antibodies were incubated for 10 minutes at 4^o^C with gentle rotation. Beads with bound antibodies were then separated from the supernatant, washed once with 200 μL of the PBS with 0.05% Tween-20 solution and stored on ice. Pre-cleaned lysate was separated from the beads used for pre-incubation and added to the beads coated with antibodies. Mixture containing lysate and magnetic beads with antibodies was incubated at 4^o^C for 20 minutes with mixing on a rotator. Beads were then collected and washed as described earlier (7). Elution was performed twice for 15 minutes in 50 μl at 65^o^C in thermoblock with shaking (1400 rpm). All 100 μl of obtained eluates were incubated with 1 μl of RNase A (100 mg/ml, Qiagen, 19101) for 30 min at 65^o^C. Then, 5 μl of Proteinase K (20 mg/ml, Qiagen, 19133) was added and the samples were incubated for 1 h at 50^o^C followed by overnight incubation at 65^o^C. Such prepared probes were stored on ice and 3 μl of 3 M sodium acetate (pH=5) was added. DNA purification was performed using the Qiaquick Qiagen PCR purification Kit according to the manufacturer’s instructions. The DNA was stored at -20^o^C. Purified DNA from two ChIP reactions with the use of 3 μl and 6 μl of antibodies was sequenced (ChIP-seq). Purified DNA from ChIP performed with empty vector strain was also sequenced and used as controls.

Sequencing of ChIP samples was performed in the Laboratory of DNA Sequencing and Oligonucleotide Synthesis of Institute of Biochemistry and Biophysics Polish Academy of Sciences in Warsaw. NGS library was constructed using the QiaSeq Ultralow Input Library kit. The sample was further quality checked on 1% agarose gel and concentration was measured using qPCR KAPA Library Quantification Kit. Libraries were sequenced using standard Illumina protocols.

Reads were quality-checked and filtered using FASTP version 0.20.0 (<https://github.com/OpenGene/fastp>) (2). Reads were mapped to *P*. *aeruginosa* PAO11161 genome (CP032126.1) using Bowtie2 version 2.3.4.3 (3) using default settings. Obtained *.sam files were sorted (samtools sort -n), run through samtools fixmate with the -m option, again sorted (samtools sort) and duplicates were marked with samtools markdup. Samtools ver. 1.9 was used (8). The files were indexed and used to generate coverage *.bigwig files, normalized to 1× sequencing depth (RPGC), without binning and smoothing using the bamCoverage tool ver 3.3.0 included in deepTools (9). Custom R script was used to import and visualize the coverage data.

BsrA peaks were identified by comparing combined replicates of IP samples (BsrA-F) for each strain with negative control (empty vector EV-F) samples using the callpeak function of MACS2 ver 2.1.2 (10) with default options for paired-end BAM files, and 0.05 as false discovery rate (FDR) cut-off. The DNA binding motif was identified using MEME version 5.3.3 (11), with any number of repetitions option, using 200 bp around sixty BsrA peak summits with the highest fold enrichment (FE ≥ 7) including summit of peak number 188 corresponding to the promoter region of *mqoA* (*PA3452*).

Raw data are available in the NCBI’s Gene Expression Omnibus (GEO) database (http://www.ncbi.nlm.nih.gov/geo/) under accession number [GSE16323](https://www.ncbi.nlm.nih.gov/geo/query/acc.cgi?acc=GSE163233)3.

*In vitro* protein-DNA interactions

DNA fragments amplified with the use of #8/#9 primers for *bsrA* promoter, #10/#11 for *mqoA* (*PA3452*) promoter and #12/#13 for *PA1112.1* promoter on the template of pMEB47, pMEB187 and/or genomic DNA were purified (Table S2). EMSA was performed in the presence of non-specific dsDNA amplified with the use of #14 and #15 primers on the template of pCM132 as a competitor in the reaction. DNA fragments (~100 nmol) were incubated in binding buffer (10 mM Tris-HCl pH 8.5, 10 mM MgCl2, 100 mM KCl, 0.1 mg/ml bovine serum albumin) with the increasing concentration of purified His_6_-BsrA in RT for 20 minutes. All reactions were prepared in a total volume of 20 μl. The samples were separated on 1.5% agarose gels in 0.5 x Tris-borate-EDTA (TBE) buffer. DNA bands were stained with 0.5 µg/ml ethidium bromide followed by destaining in water and visualized on a UV transilluminator. The same *PA1112.1* promoter DNA was also used to obtain its shortened version by digestion of *PA1112.1*p with SspI restriction enzyme cutting off its part (71 bps) followed by purifying of the 235 bps DNA fragment containing promoter lacking T-N_11_-A motif.

Regulatory experiments with promoter-*xylE* fusions in *E. coli*

*E*. *coli* DH5α double transformants carrying pPTO1 derivatives with *xylE* reporter gene and pAMB9.37 (*lacI^Q^*-*tac*p) derivatives encoding tested proteins were used to perform catechol 2, 3-oxygenase activity (the product of *xylE*) assay. The following variants:

pMEB190 (*bsrA*p-*xylE*)/pAMB9.37 (*lacI^Q^*-*tac*p), pMEB190 (*bsrA*p-*xylE*)/pMEB63 (*lacI^Q^*-*tac*p-*BsrA*), pMEB232 (*pa1112.1*p-*xylE*)/pAMB9.37 (*lacI^Q^*-*tac*p), pMEB232 (*pa1112.1*p-*xylE*)/pMEB63 (*lacI^Q^*-*tac*p-*BsrA*) were tested to observe influence of BsrA expression on the activity of indicated promoters. Transformants carrying the following pairs of plasmids: pMEB190 (*bsrA*p-*xylE*)/pAMB9.37 (*lacI^Q^*-*tac*p), pMEB190 (*bsrA*p-*xylE*)/pMEB230 (*lacI^Q^*-*tac*p-*PA2551*), pMEB190 (*bsrA*p-*xylE*)/pMEB231 (*lacI^Q^*-*tac*p-*PA3587*), pMEB190 (*bsrA*p-*xylE*)/pMEB236 (*lacI^Q^*-*tac*p-*PA3398*), pMEB190 (*bsrA*p-*xylE*)/pMEB237 (*lacI^Q^*-*tac*p-*PA4902*) and pMEB190 (*bsrA*p-*xylE*)/pMEB238 (*lacI^Q^*-*tac*p-*PA5189*) were used to test impact of different transcriptional regulators on *bsrA* promoter activity. In every variant of experiment, cells with the promoterless pPTOI (-*xylE*) and pAMB9.37 (*lacI^Q^*-*tac*p) were used as a background control.

Cell extracts were prepared from exponentially growing cultures grown at 37^o^C in LB medium supplemented with 50 µg/ml kanamycin and 10 µg/ml chloramphenicol (in the case of control empty pPTO1 only 50 µg/ml kanamycin). The assay was performed as previously described (12). Obtained measurements of XylE activity were normalized to the amount of the protein in extracts indicated by Bradford assay (13). Data represent mean values from three measurements ±SD.

Tests of kanamycin efficacy

Stationary-phase cells (after overnight incubation at 37^o^C with shaking) of WT *P. aeruginosa* PAO1161 and PAO1161 Δ*bsrA* strain were harvested by centrifugation, washed once in 1 mL of PBS buffer, and re-suspended in 1 mL of M9 medium containing 50 µg/ml kanamycin with appropriate carbon source addition - 30 mM sodium acetate, 15 mM fumarate, 20 mM PEP (phosphoenolpyruvate). The concentration of compounds used for carbon supplementation was normalized to achieve a 60 mM final carbon concentration. 50 µl of each sample was collected in sterile Eppendorf tube for cfu enumeration in the starting point and then samples were incubated at 37^o^C with shaking for 4 hours. After that time, the 50 µl of each sample was collected. Dilutions (in the range from 10^0^ to 10^-7^) in the amount of 20 µl, from t_0_ and t_4_, were plated on LB agar. After overnight incubation of plates at 37^o^C, the number of colonies was assessed.

DNA pull-down assay

Biotinylated fragments of DNA containing the *bsrA* promoter region were amplified by PCR using #8 and #16 primers (Table S2) and pMEB47 plasmid DNA as a template. As the reaction specificity control, a biotinylated fragment of empty pCM132 vector obtained with the use of #16 and #17 primers was also prepared. DNA was purified by gel-extraction and about 10 µg of specific and non-specific DNA was used in a single reaction with 100 μl of magnetic beads coated with streptavidin (Dynabeads M-280 Streptavidin, Invitrogen, 11205D). Cell extracts from cultures in the exponential phase of *P. aeruginosa* PAO1161 (WT) strain and from overproducing strain carrying pMEB99 vector were used for incubation with magnetic beads coupled with DNA fragments. Following probes were prepared to combine beads with DNA and extract: WT + specific DNA; WT + non-specific DNA; BsrA-FLAG overproducer + specific DNA. The elution of proteins attached to the beads was performed in the buffer with increasing NaCl concentrations (0.2 M, 0.5 M, 1 M). The eluates from the variant prepared with BsrA-FLAG overproducing strain were tested by SDS-PAGE followed by Western blot with anti-FLAG antibodies as a confirmation of predicted binding of BsrA-FLAG protein to *bsrA*p and correctness of the pull-down procedure. The probes from WT strain and specific or non-specific DNA were examined by SDS-PAGE. Proteins from selected fractions (from second and third elution) were identified by mass spectrometry analysis (Mass Spectrometry Laboratory IBB PAS). Results of the analysis were compared to each other to indicate proteins which specifically attached to the *bsrA*p promoter.
