## Supplementary figures and images for "The LysR-type transcriptional regulator BsrA (PA2121) controls vital metabolic pathways in *Pseudomonas aeruginosa*"

### Figure S1

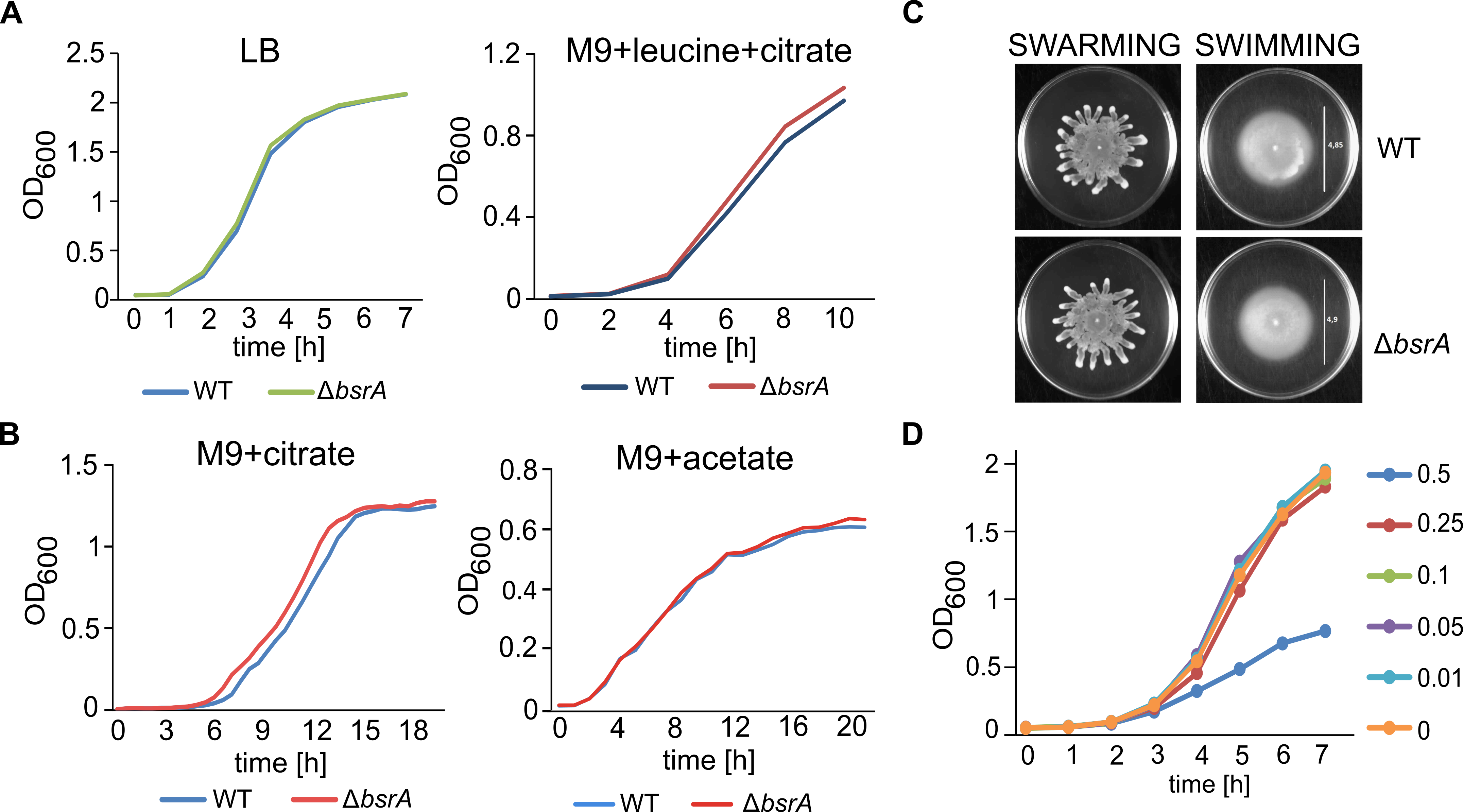

### Figure S2

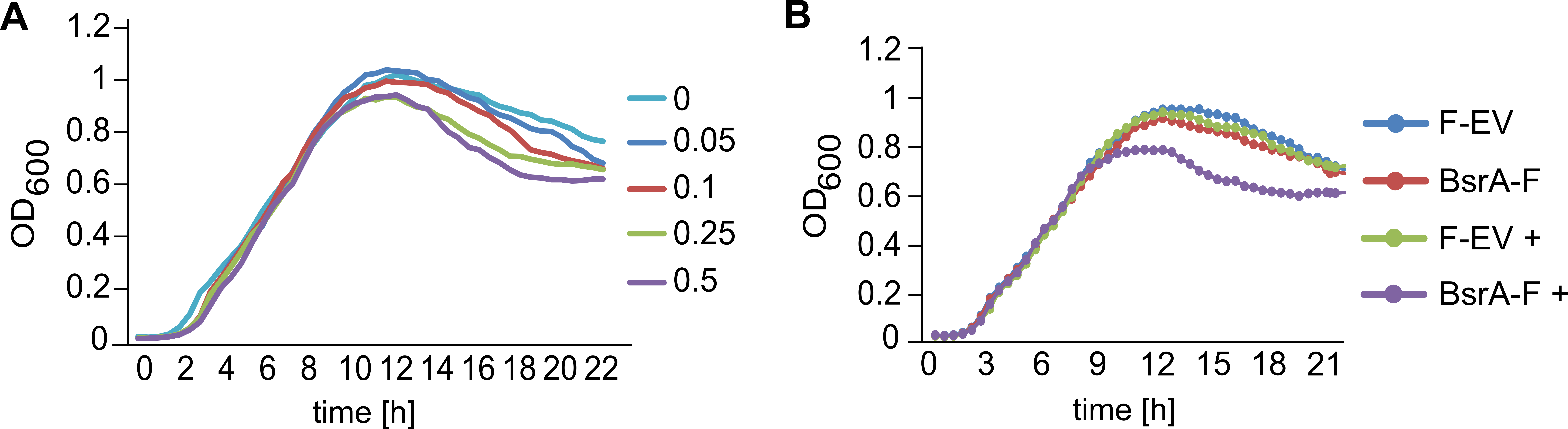
